# Single-Nucleus Analysis of Human White Adipose Tissue Reveals Adipocyte Subsets with Distinct Metabolic Profiles

**DOI:** 10.1101/2025.09.14.673351

**Authors:** Vissarion Efthymiou, Adhideb Ghosh, Sean D. Kodani, Xavier Caubit, Laurent Fasano, Waqar Ali, Lindsay S. Poulos, Henrique Camara, Anushka Gupta, Yasmine Belaidouni, A. Sina Booeshaghi, Shiyi Yang, Ruchir Rastogi, Farnaz Shamsi, Ashley Vernon, Aaron Streets, Yu-Hua Tseng, Mary Elizabeth Patti

## Abstract

Anatomic location of white adipose tissue is a determinant of cardiometabolic risk. To understand differences within/between adipose depots, we generated 65,668 single-nucleus transcriptomes from human subcutaneous or intraabdominal adipose tissue (SAT/IAT). Unsupervised analysis revealed 26 adipose-resident cell clusters including two subpopulations of mature adipocytes, characterized by high vs. low expression of adipocyte maturation genes (ADIPOMAT^hi^ vs. ADIPOMAT^lo^). ADIPOMAT^lo^ adipocytes demonstrate a low-differentiation, pro-inflammatory, and pro-fibrotic transcriptome. IAT-resident ADIPOMAT^lo^ were more abundant in higher BMI donors, while SAT-resident ADIPOMAT^lo^ associated with impaired glycemia. TSHZ3 was identified as a candidate regulator of ADIPOMAT^lo^ transcriptome. TSHZ3 knockdown in adipogenic progenitors inhibited differentiation, with downregulation of early adipogenic regulators (e.g. CEBPA/B, PPARG) and mature adipocyte genes. Heterozygous deletion of *Tshz3* in mice reduced SAT and IAT weight. Here, we show that adipocyte subsets with distinct transcriptomic signature reside in human WAT; altered TSHZ3-mediated transcriptional regulation may contribute to low-maturation subpopulation linked to metabolic disease.

## Introduction

The rising epidemic of obesity poses a severe threat to personal and public health worldwide. Obesity is associated with multiple cardiometabolic diseases, including type 2 diabetes (T2D)^1^, metabolic fatty liver disease (MAFLD)^2^, cardiovascular disease^3^, and multiple types of cancer^4^. Currently, anti-obesity therapies include caloric restriction, bariatric surgery, and increasingly potent incretin analogues^5^, but all are complicated by side effects, cost, and uncertain durability. Moreover, no effective approaches to specifically reduce development and expansion of metabolically unhealthy adipose tissue or to promote healthy expansion are available. Thus, a deeper molecular understanding of the mechanisms promoting adipose health - and how these are dysregulated in metabolically unhealthy obesity - is required.

Adipose tissue is highly heterogeneous, with both lipid-laden adipocytes and a diverse stromal vascular fraction (SVF) containing adipogenic progenitor cells (APC), endothelial cells, smooth muscle cells (SMC), macrophages and other immune cells. While adipocytes comprise the majority of adipose tissue by volume, the majority of cells - by number - are within the SVF. Lineage-tracing and cell-sorting studies have demonstrated that hyperplastic adipose expansion relies on differentiation of APCs in the SVF^6,7^, broadly marked by expression of PDGFRα^8–10^.

Perturbations in adipocyte mass, differentiation, and function are intimately linked to metabolic health. Both increases in adipose mass, as with obesity, and reductions in mass, as with lipodystrophy, are associated with insulin resistance and risk for type 2 diabetes. In both situations, the functional capacity of adipocytes to store lipid appropriately is exceeded; this leads to pro-inflammatory signaling and ectopic accumulation of lipid in key metabolic tissues, contributing to insulin resistance and reduced insulin secretion^11^. Additional adipose-resident cells, such as immune cells, may also contribute to altered metabolic function of adipose tissue in this setting. Adipose tissue macrophages (ATMs) may acquire a classically activated pro-inflammatory (M1) or an alternatively activated anti-inflammatory (M2) phenotype, which play important role in adipose metabolic health. This distinction has been lately challenged as simplistic due to the large heterogeneity of macrophages revealed via single-cell studies^12^. For instance, obesity leads to accumulation of lipid-associated macrophages (LAMs), which are distinct from the classically activated M1 ATMs^13^.

With the advancement of single-cell and single-nuclei RNA sequencing (RNA-seq) technologies, multiple novel cell populations with stem, adipogenic, or anti-adipogenic potential and other previously unknown functions have been revealed within both white and brown adipose tissue^14,15^. Thus, a key scientific goal with implications for novel therapeutics is to identify how cell populations and their transcriptomes differ in metabolic disease.

In this work, we report characterization of the nuclear transcriptome derived from human IAT and SAT biopsies. Here, we identify a metabolically distinct subtype of adipocytes based on low expression of markers of adipocyte maturation, demonstrate differential abundance of this population in relation to BMI and glycaemia, and identify TSHZ3 as a novel upstream regulator of adipogenic differentiation.

## Results

### Single-nucleus RNA-Sequencing (snRNA-Seq) reveals 26 distinct adipose-resident cell clusters with differing distribution in SAT and IAT

Nuclei were isolated for single-nucleus RNA-sequencing (snRNA-Seq) (**Figure 1A**) from 22 human SAT and IAT biopsies obtained during elective abdominal surgery; 6 were paired biopsies (SAT and IAT from the same donor). Intraabdominal samples were derived from both gastrocolic (n=9) and mesenteric (n=3) locations. Mean age of the 19 donors was 41±3 years, with body mass index (BMI) ranging from 23.6 to 60.9 kg/m^2^ (**Figure 1B).** snRNA-Seq libraries were processed to remove doublets, low quality cell barcodes, and background ambient RNA (**Figure S1A**); all samples were integrated using scVI-tools (see Methods)^16,17^. A total of 65,668 high-quality nuclei were distributed across multiple cell clusters in all biopsy samples (**Figure S1B-C**). Unsupervised hierarchical clustering grouped cells into 26 distinct clusters (**Figure 1C and S1D).**

**Figure 1.**
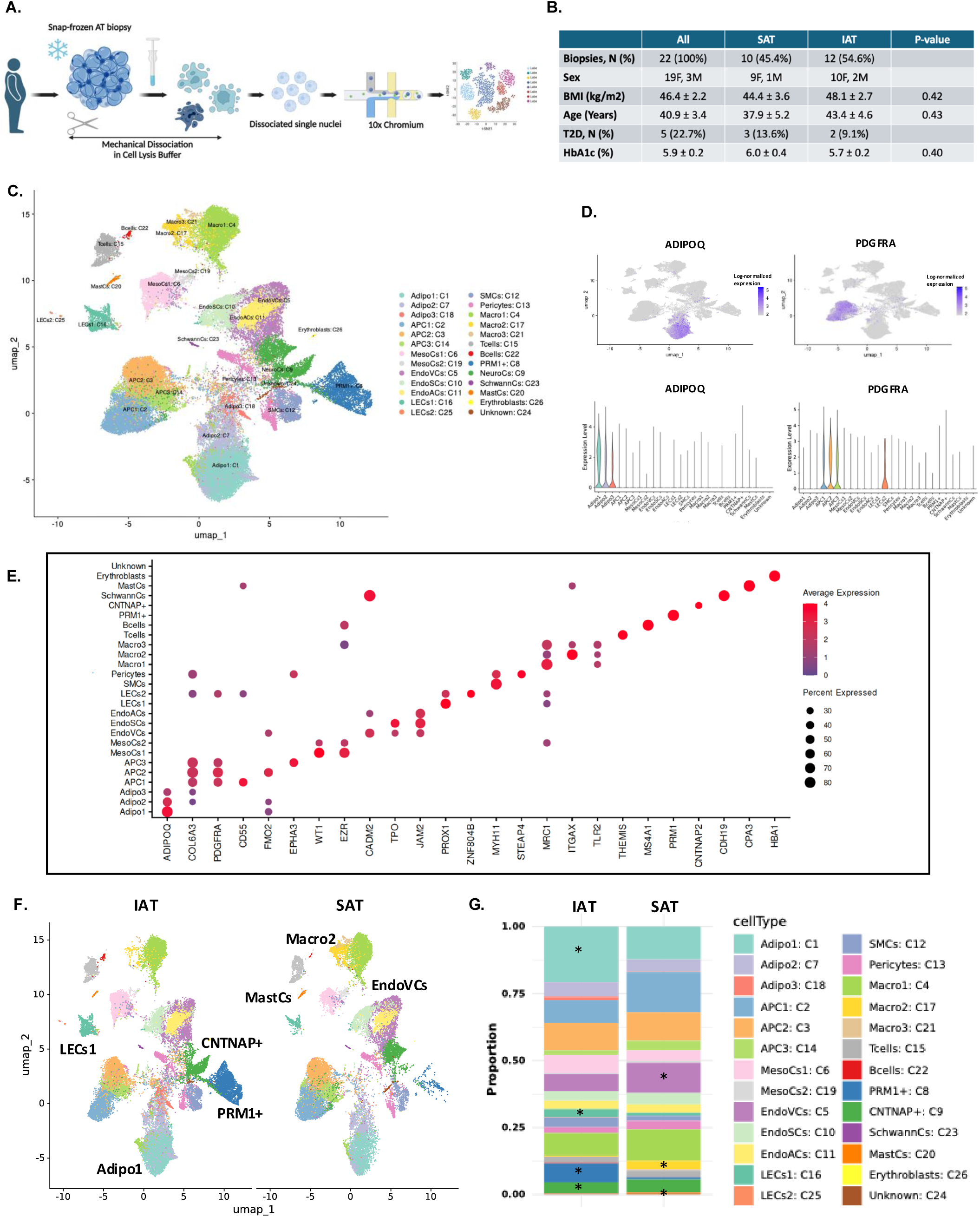
Human abdominal subcutaneous (SAT) and intraabdominal (IAT) white adipose tissue (WAT) is highly heterogeneous, with depot-specific differences in cell types. A. Workflow to acquire single-nuclei RNA-Seq data from frozen WAT biopsies, B. Participant demographics, C. UMAP Plot showing cell clusters identified during unsupervised analysis, with identification of cell types, based on cell type calling analyses, D. Single-nucleus gene expression (upper: Feature Plot, lower: Violin Plot) of ADIPOQ and PDGFRA in all captured nuclei, E. DotPlot showing expression of top cell type-specific markers for each identified cell cluster, F. UMAP Plot showing all identified cell clusters divided by anatomical location (IAT: left, SAT: right). Labeled cell types indicate enrichment of that particular cell type in the indicated depot, G. Quantification of percentage of cell types identified in IAT and SAT biopsies. * indicates p<0.05 for difference between IAT vs. SAT.

To identify cell types associated with each cluster, we employed two complementary methods (detailed in Methods). First, we identified markers for each cluster using an unsupervised approach using Seurat (**Figure S1E**). Secondly, we used established markers curated from the literature and performed reference mapping to published data from Emont et al^15^ (**Figure S1F**), to predict cell types in a supervised “cell-calling” manner. For example, cell populations with significant expression of adiponectin (ADIPOQ) are annotated as mature adipocytes, whereas those with robust PDGFRA expression are annotated as adipogenic progenitor cells (APC) (**Figure 1D**). Additional clusters (**Figure 1C**) included mature adipocytes (Adipo, clusters 1, 7, 18), APC (2, 3, 14), smooth-muscle cells (SMC, 12), pericytes (13), endothelial cells (endothelial venular or EndoVCs, endothelial stalk or EndoSCs, and endothelial arteriolar or EndoACs, 5, 10, 11, respectively), erythroblasts (26), lymphatic endothelial cells (LEC, 16, 25), mesothelial cells (MesoCs, 6, 19), Schwann cells (23) and immune cells, including T cells (15), B cells (22), mast cells (20), and subpopulations of macrophages (Macro, 4, 17, 21). The top enriched markers for each cluster relative to all others are provided in **Supplementary Table 1,** and in a dot plot (**Figure 1E)** and heat map **(Figure S1E)**. We also identified two novel clusters: one with neuronal ontology was enriched for multiple members of the contactin-associated protein-like family (CNTNAP), thus termed neuronal-associated cells (NeuroCs, cluster 9); and one with protamine 1 (PRM1) as its top marker, termed PRM1+ cells (cluster 8) (**Figure 1C and Figure S1G-H)**.

Cells within each cluster were observed in both SAT and IAT samples (**Figure 1F**, **1G, and S1I**), and across the range of age and BMI. However, the distribution of cell types, expressed as a percentage of all cells, differed between IAT and SAT (**Figure 1F**, **1G, and S1I**); there were few differences between gastrocolic and mesentery IAT (**Figure S1C**). Specifically, the percentage of mature adipocytes (Adipo1, cluster 1) and lymphatic endothelial cells (LECs1, cluster 16) were significantly increased in IAT, whereas that of Endothelial Venular Cells (EndoVCs, cluster 5), Macrophages II (Macro2, cluster 17), and a low abundance population of mast cells (MastCs, cluster 20) were significantly decreased in IAT (**Figure S1I**). Moreover, both the neuronal-associated CNTNAP+ and PRM1+ populations were almost exclusively present in IAT biopsies (**Figure S1G-S1H**). To our knowledge, this is the first time that these two subtypes are identified specifically in IAT. The functional role of these cell types remains to be determined in future studies.

### Identification of APC subtypes with distinct adipogenic and inflammatory patterns and anatomical location

Using unsupervised clustering, we identified three distinct adipogenic progenitor cell (APC) populations, termed APC1 (cluster 2), APC2 (cluster 3), and APC3 (cluster 14) (**Figure 1C and S2A**), all marked by expression of PDGFRA (**Figure 1D**). As predicted, two APC clusters (APC1 and APC2) were enriched for Epithelial Mesenchymal Transition signaling pathways (**Figure S2B**); APC3 was enriched for Hedgehog and Wnt–Beta Catenin signaling pathways (**Figure S2B**).

Analysis of differentially expressed genes between these clusters is provided in **Supplementary Figure 2A** (list of APC markers in **Supplementary Table 1;** genes differentially expressed between APC clusters in **Supplementary Table 2**). Pairwise cluster analysis using ENRICHR revealed enrichment of Inflammatory Response, TGF-Beta Signaling, Androgen Response, and IL-2/STAT5 Signaling pathways in APC1, whereas Adipogenesis, Estrogen Response, and Bile Acid Metabolism were significantly enriched in APC2 (**Supplementary Table 2**). Likewise, APC3 was enriched for pro-adipogenesis pathways relative to APC1, but to a lesser degree than APC2 (**Supplementary Table 2**). Interestingly, multiple mouse adipose single-cell studies suggest there are two major adipogenic progenitor cell types, one that is pro-fibrotic with lower adipogenic capacity (potentially corresponding to APC1), and one that is pro-adipogenic (which could correspond to APC2/3 in the current data set)^18^. Reference mapping to the Emont et al^15^ dataset shows resemblance of the transcriptome of APC1 to the CD55+ fibro-adipogenic progenitors (FAPs), APC2 to FMO2+ pre-adipocytes, and APC3 to EPHA3+ adipogenesis-regulators (Aregs) cells.

Using CheA3 analyses, we predicted the top 10 putative TFs regulating depot-specific gene expression in each of the 3 APC subtypes. TFs predicted to uniquely regulate the 40 genes (log_2_FC>1, adj-p<0.05) with greater expression in IAT- vs. SAT-resident APCs included EBF1, BNC2, TCF21, TSHZ3, ZFHX4, ZNF385D, FOXP2, NFIA, PBX1, and LHX9; BNC2 was common to all 3 APC subtypes (**Figure S2C-D**). Conversely, HOXA6, members of the HOXC (HOXC5, HOXC10, HOXC11) and HOXD (HOXD4, HOXD8, HOXD9, HOXD10, HOXD11) families of TFs and TWIST2 were predicted to be regulatory for the 42 genes with greater expression in SAT-resident APCs. TBX18 and TWIST2 were common to all 3 APC subtypes (**Figure S2C-D**). Notably, these data parallel rodent studies demonstrating higher expression of Shox2, HoxC8, and HoxA5 in SAT (inguinal) vs. IAT (mesenteric) depots^19^.

### Identification of two subpopulations of mature adipocytes differing by expression of genes marking adipocyte maturity (ADIPOMAT^hi^ and ADIPOMAT^lo^)

Mature adipocytes constitute the major functional units of adipose tissue. These represented 18.5% of total nuclei in SAT and 25.0% in IAT. These percentages are consistent with other published studies evaluating cell heterogeneity in human WAT^15,20^. We identified two clusters of mature adipocytes (Adipo1 and Adipo2, or clusters 1 and 7) (**Figure 1C**) based on robust expression of multiple adipocyte markers, including ADIPOQ, PPARG, PLIN, and LEP, and on reference mapping to published single-nucleus adipose datasets. Using the analytical tool VISION^21,22^ to plot gene expression signatures (instead of individual genes) at a single-nucleus level, we confirmed that both of these clusters have greater enrichment of adipogenesis signatures^23^ than all other clusters (**Figure 2A**). However, adipogenesis signatures were significantly greater in Adipo1 than Adipo2; given that collective expression of these genes defines maturity of adipocytes, we termed these populations adipocyte-maturation-high (ADIPOMAT^hi^) (Cluster 1) and adipocyte-maturation-low (ADIPOMAT^lo^) (Cluster 7) adipocytes; these patterns were observed whether expressed per cell or per biopsy (**Figure 2B**). In agreement, reduction in adipogenesis scores in the ADIPOMAT^lo^ population was significant in 21 of 22 biopsy samples (adj-p<0.05, **Figure S2E**). These data are broadly concordant with the concept that some human adipogenic progenitors are highly committed toward adipogenesis but express low levels of adiponectin^24^. We hypothesized that ADIPOMAT^hi^ were likely to represent canonical mature adipocytes, whereas ADIPOMAT^lo^ adipocytes could represent an alternative adipocyte type or state with incomplete maturation or with partial de-differentiation. Transcript patterns within these clusters differed (**Figure 2C**), with greater expression of mature adipocyte markers in the ADIPOMAT^hi^ cluster 1 (**Figure 2D**). Genes upregulated in ADIPOMAThi adipocytes which are involved in enhanced adipogenesis and fatty acid metabolism include ACACB, ACADM, ACADL, ACOX1, ACDLY6, and CD36 (**Figure S2F**), whereas genes upregulated in ADIPOMATlo adipocytes within pro-inflammatory and pro-fibrotic pathways include COL15A1, COL6A3, IL6ST, PDLIM5, RABGAP1L, and LEPR (**Figure S2G**). Notably, a transcriptomic signature of ADIPOMAT^lo^ adipocytes aligns with “non-canonical” ECM-secreting pro-fibrotic adipocyte subtypes (**Figure S2H**) that were recently captured in human abdominal subcutaneous and visceral WAT biopsies^25^. Markers of these “non-canonical” adipocytes, including PTRB, PDE4D, and SKAP1 also have increased expression in ADIPOMAT^lo^ vs. ADIPOMAT^hi^ populations in our datasets, and their expression in human adipocytes has been validated at the protein level^25^ (**Figure S2I**).

**Figure 2.**
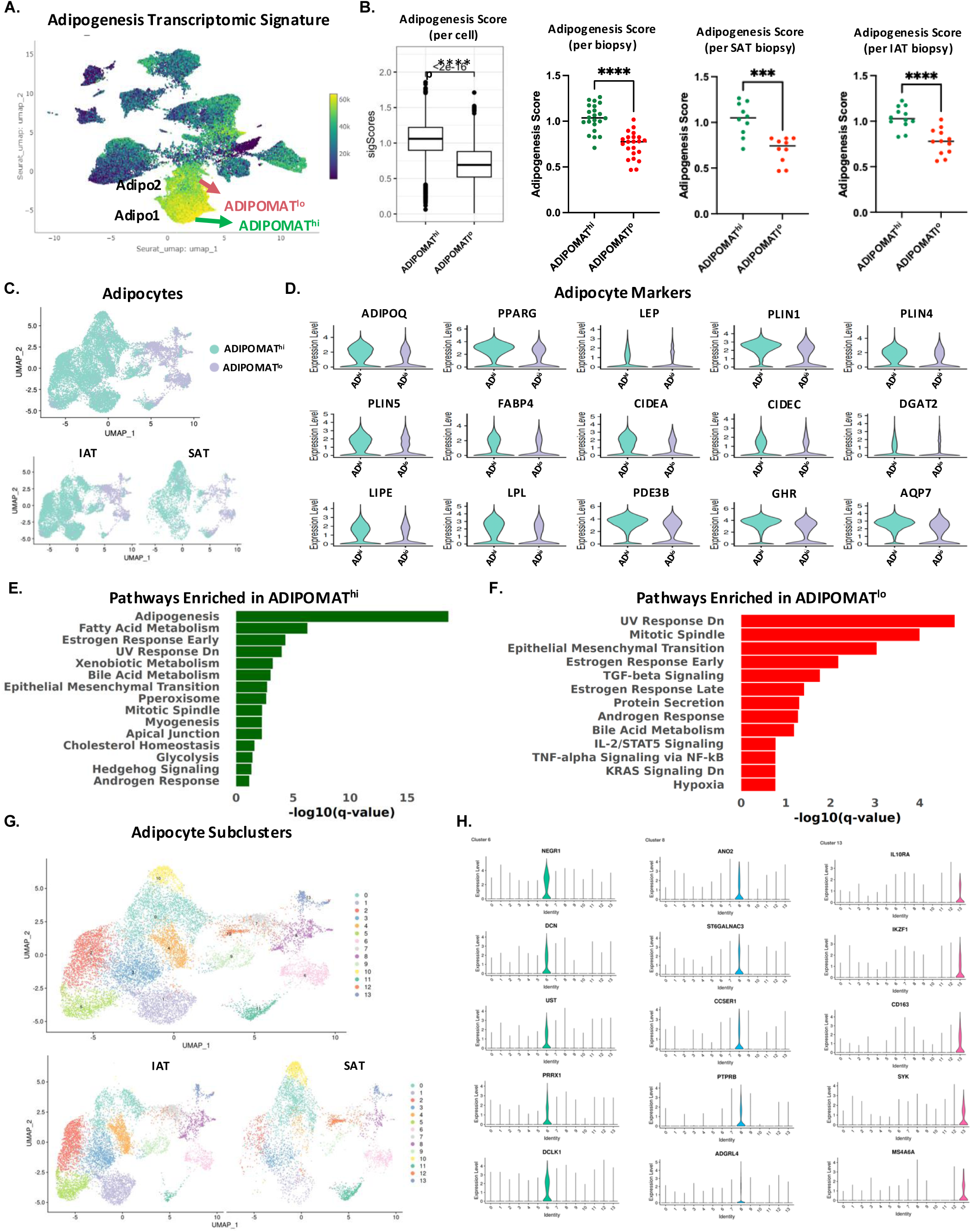
Identification of two distinct mature adipocyte subpopulations, termed ADIPOMAT^hi^ and ADIPOMAT^lo^, with ADIPOMAT^lo^ adipocytes having a less adipogenic and more pro-inflammatory transcriptomic signature. A. UMAP plot colored by adipogenesis transcriptomic signature, calculated in VISION. B. Adipogenesis transcriptomic signature score in ADIPOMAT^hi^ and ADIPOMAT^lo^ adipocytes. Score was calculated per cell (left panel) per biopsy, or per biopsy within SAT or IAT***p<0.001, ****p<0.0001. C. UMAP Plot showing re-clustered mature adipocytes (ADIPOMAT^hi^ and ADIPOMAT^lo^) for all biopsies (upper) or divided by anatomical location (IAT: left, SAT: right) (lower). D. Violin plots showing expression of established mature adipocyte markers in ADIPOMAT^hi^ and ADIPOMAT^lo^ cell clusters. E. Pathway enrichment ontology analysis for genes upregulated in ADIPOMAT^hi^ vs. ADIPOMAT^lo^. F. Pathway enrichment ontology analysis for genes upregulated in ADIPOMAT^lo^ vs. ADIPOMAT^hi^. For both E and F, input genes had adjusted-p<0.05, log_2_ fold-change>0.5. G. UMAP Plot showing subpopulations of re-clustered mature adipocytes (ADIPOMAT^hi^ and ADIPOMAT^lo^) for all biopsies (upper) or divided by anatomical location (IAT: left, SAT: right) (lower). H. Violin Plots showing the top 5 distinct markers of ADIPOMAT^lo^ adipocyte subpopulations 6 (left), 8 (middle), and 13 (right).

To determine whether the anatomic localization of the two mature adipocyte subclusters influenced their transcript patterns, we analyzed the transcriptome of the ADIPOMAT^hi^ and ADIPOMAT^lo^ adipocytes in SAT vs. IAT depots, using Seurat differential expression (DE) analysis (see Methods). Within ADIPOMAT^hi^, 1,039 genes were upregulated in SAT and 668 genes upregulated in IAT; within ADIPOMAT^lo^, 660 genes were upregulated in SAT and 547 genes upregulated in IAT (log_2_FC>0.5, adj-p<0.05). ENRICHR pathway analysis revealed similarities in the depot-specific transcriptome of adipocyte sub-types: epithelial-mesenchymal transition ontology was consistently enriched in genes upregulated in SAT vs. IAT in both ADIPOMAT^hi^ and ADIPOMAT^lo^ clusters (**Figure S2J and Supplementary Table 2**). Conversely, there was enrichment of UV, Androgen, and Estrogen Response, and TGF-beta Signaling within genes upregulated in IAT (**Figure S2J**). Differential gene expression analysis of ADIPOMAT^lo^ vs. ADIPOMAT^hi^ showed increased expression of COL6A3 in ADIPOMAT^lo^ (both in IAT and SAT), while expression of LEPR was increased in ADIPOMAT^lo^ vs. ADIPOMAT^hi^ within SAT biopsies (**Figure S2K**). CheA3 analyses to predict regulatory TFs of the transcriptome of the two mature adipocyte subtypes across depots (**Figure S2L**) revealed FOXO1 among the predicted regulators in IAT; FOXO1 has been previously shown to impact adipogenesis specifically in visceral adipose^26^. Likewise, SAT-enriched transcriptional regulators included PRRX1, TBX18, and TWIST2 (**Figure S2M**).

### Transcriptome of ADIPOMAT^lo^ adipocytes is enriched for genes associated with impaired function of adipose tissue

Differential expression analysis of ADIPOMAT^hi^ vs. ADIPOMAT^lo^ adipocytes (combined for both IAT and SAT) using Seurat (see Methods) identified 525 significantly differentially expressed transcripts (adj-p<0.05, log_2_FC>0.5), with 75 increased in ADIPOMAT^lo^ and 450 upregulated in ADIPOMAT^hi^ (**Supplementary Table 2**). ENRICHR Pathway analysis revealed 20 Hallmark gene sets enriched in transcripts upregulated in ADIPOMAT^hi^ (q<0.05), including adipogenesis and fatty acid metabolism pathways (**Figure 2E**); conversely, there was enrichment of inflammatory pathways, including TGF-β, IL2/STAT5, and TNF-α via NF-κB signaling pathways in transcripts upregulated in ADIPOMAT^lo^ (**Figure 2F**).

Sub-clustering of the ADIPOMAT^hi^ and ADIPOMAT^lo^ mature adipocyte populations revealed transcriptional heterogeneity. 14 different subtypes were identified, including three distinct subpopulations of ADIPOMAT^lo^ adipocytes (subclusters 6, 8, and 13), and one subpopulation of EBF2+/PRDM16+ “beige” adipocytes (subpopulation 5) (**Figure 2G and S2N-O)**. The EBF2+ population represents thermogenic beige adipocytes, as evidence by enriched expression of PPARGC1A and PRDM16 (**Figure S2N**). Interestingly, the abundance of beige adipocytes is approximately 3-fold higher (p<0.05) in IAT vs. SAT (**Figure S2O**). In addition, there was a strong trend for reduction in beige adipocyte (subcluster 5) abundance (p=0.07) in biopsies from patients with higher HbA1c (**Figure S2O**). Distinct markers for ADIPOMAT^lo^ subpopulation 6 included ANK2, DCN, NEGR1, COL6A3, PRRX1, and DCLK1; for ADIPOMAT^lo^ subpopulation 8 included ELMO1, MECOM, PTPRB, and ANO2; and for ADIPOMAT^lo^ subpopulation 13 included IL10RA, SYK, MERTK, PDE4D, and PTPRC (**Figures 2H, S2P-Q**). As shown before, the presence of these markers in mature adipocytes has been validated in an independent cohort via immunohistochemistry (**Figure S2I**).

Together, these findings suggest that ADIPOMAT^hi^ adipocytes represent a more functional mature adipocyte subtype, which is actively involved in lipid metabolism, whereas ADIPOMAT^lo^ adipocytes represent a more pro-inflammatory mature adipocyte subtype with attenuated metabolic function.

### Mature adipocytes, and specifically ADIPOMAT^lo^ adipocytes, are more prevalent in participants with T2D and correlate with body mass index (BMI) and glycaemia (HbA1c)

Next, we used linear regression (correlation) analysis to examine the relative contribution of mature adipocytes, and ADIPOMAT^hi^ and ADIPOMAT^lo^ sub-populations, within SAT and IAT and their relation to metabolic phenotypes of the biopsy donors. The percentage of mature adipocytes within the total population of cells captured was significantly higher in IAT as compared to SAT (**Figure 1G and S1G**). However, there was no significant difference in the percentage of ADIPOMAT^hi^ and ADIPOMAT^lo^ adipocytes within IAT vs. SAT depots, whether expressed as percentage of total cells or total adipocytes (**Figure 3A**).

**Figure 3.**
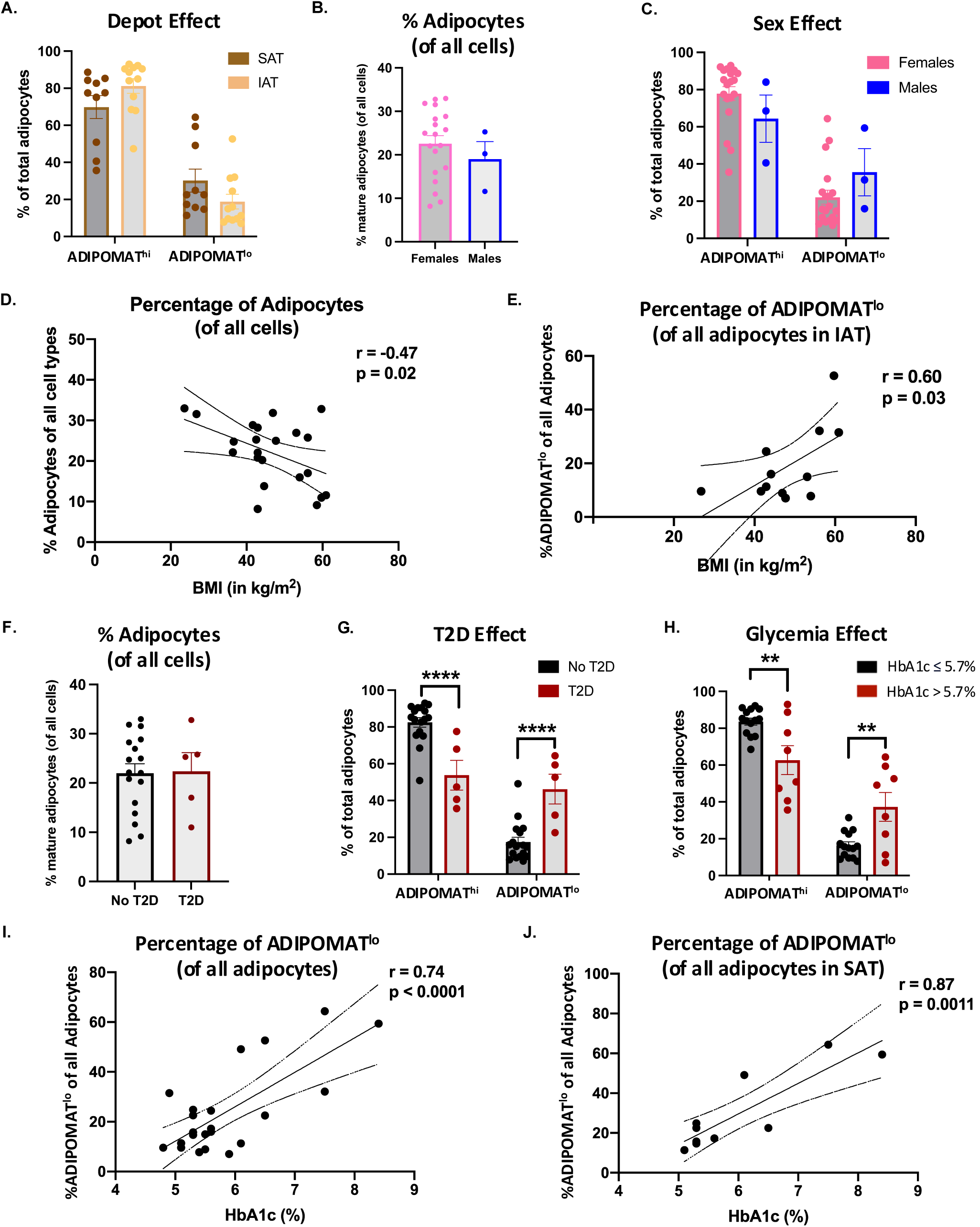
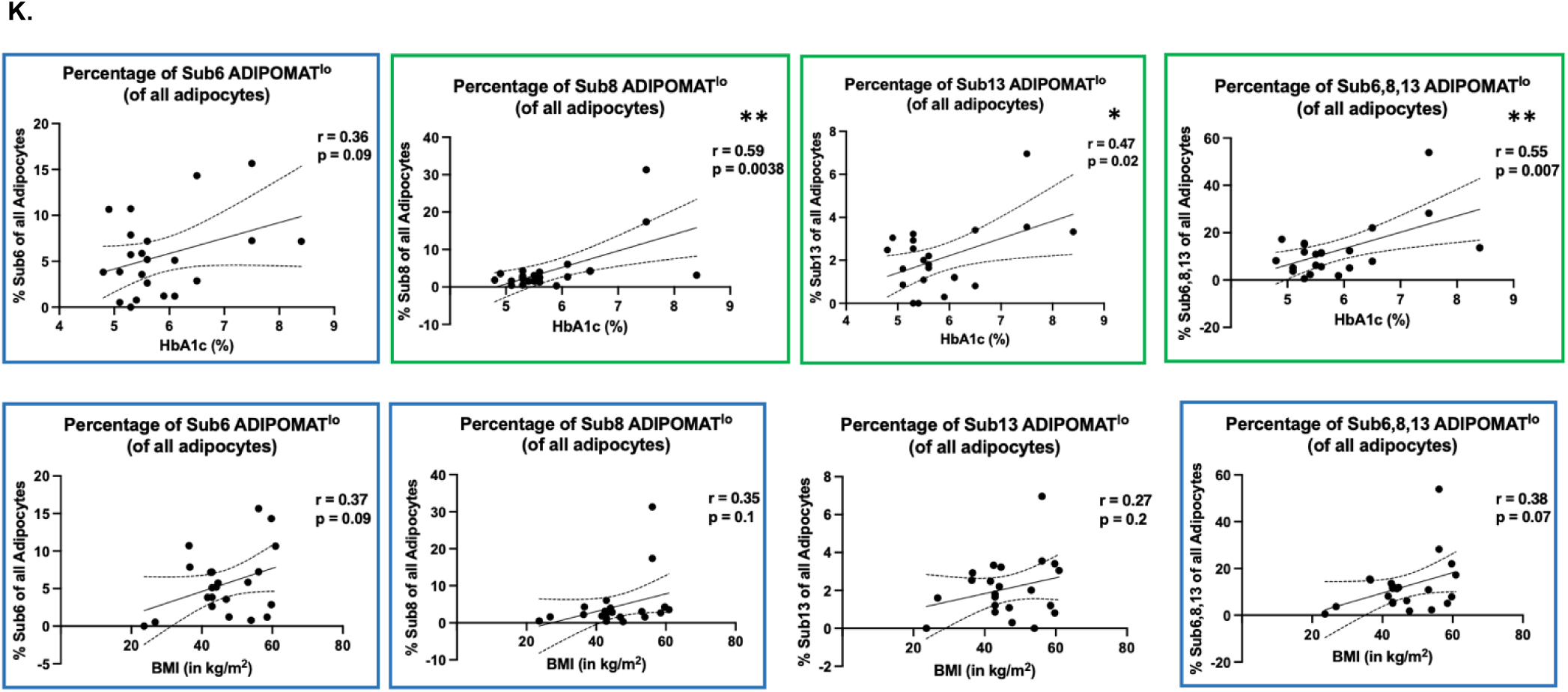
Relative abundance of ADIPOMAT^lo^ adipocytes is positively correlated with adiposity and glycaemia. A. Relative abundance of ADIPOMAT^hi^ and ADIPOMAT^lo^ adipocytes (expressed as % of all mature adipocytes) within each depot. B. Percentage of mature adipocytes (expressed as % of all cells) for male and female donors. C. Percentage of ADIPOMAT^hi^ and ADIPOMAT^lo^ (expressed as % of all mature adipocytes) for male and female donors. D. Correlation analysis of percentage of adipocytes (expressed as % of all cells) and donor BMI. E. Correlation analysis of percentage of ADIPOMAT^lo^ (expressed as % of all mature adipocytes) in IAT biopsies and donor BMI. F. Percentage of mature adipocytes (expressed as % of all cells) shown by presence or absence of T2D in donors. G. Percentage of ADIPOMAT^hi^ and ADIPOMAT^lo^ adipocytes (expressed as % of all mature adipocytes) by presence or absence of T2D in donors. H. Percentage of ADIPOMAT^hi^ and ADIPOMAT^lo^ adipocytes (expressed as % of all mature adipocytes), stratified by donor HbA1c levels. Graphs are presented as mean +/− SEM. **p<0.01, ****p<0.0001. I. Correlation analysis of percentage of ADIPOMAT^lo^ (expressed as % of all mature adipocytes) and HbA1c (%). J. Correlation analysis of percentage of ADIPOMAT^lo^ (expressed as % of all mature adipocytes) in SAT biopsies with HbA1c (%). K. Correlation analyses of percentages of ADIPOMAT^lo^ subpopulations 6, 8, 13, and 6-8-13 combined (expressed as % of all mature adipocytes) with HbA1c (%) (upper panels) and with BMI (lower panels).

Age of participants was not significantly associated with abundance (percentage) of either mature adipocytes or ADIPOMAT^hi^ or ADIPOMAT^lo^ subtypes (**Figure S3A-S3D**). Likewise, there was no significant difference in the percentage of mature adipocytes in biopsies obtained from male vs. female donors (**Figure 3B**). However, there was a trend for higher percentage of ADIPOMAT^lo^ adipocytes in male-derived biopsies (p=0.2, **Figure 3C**).

There was a significant inverse correlation between BMI and the percentage of mature adipocytes within both depots (r= −0.47, p=0.02, **Figure 3D**); this relationship was even stronger in SAT (r= −0.79, p=0.007) (**Figure S3E**), with a similar but non-significant pattern in IAT (r= −0.39, p=0.19) (**Figure S3F**). By contrast, the percentage of ADIPOMAT^lo^ adipocytes was positively correlated with BMI for IAT biopsies (r=0.60, p =0.03, **Figure 3E**), with similar trend for all biopsies (r=0.39, p=0.07) (**Figure S3G**). These relationships were corroborated using VISION. BMI was inversely associated with adipogenesis signatures (r=-0.37, p=0.02, **Figure S3H**) and positively associated with inflammatory and pro-fibrotic signatures (r=0.32, p=0.1, **Figure S3I**), with greater magnitude for ADIPOMAT^lo^ nuclei, specifically those in IAT. Thus, ADIPOMAT^lo^ adipocytes have lower adipogenesis-related and enhanced inflammation-related gene expression patterns, particularly with increasing BMI of donor.

We did not observe a significant difference in the percentage of total mature adipocytes in adipose tissue derived from individuals with vs. without T2D or association with HbA1c (**Figure 3F**, **Figure S3J**). However, there was a significantly higher percentage of ADIPOMAT^lo^ adipocytes in individuals with T2D vs. those without T2D (**Figure 3G**). Likewise, the percentage of ADIPOMAT^lo^ adipocytes was greater in those with higher HbA1c (**Figure 3H**), with the abundance of ADIPOMAT^lo^ adipocytes showing a significant positive correlation with increasing HbA1c (r=0.74, p<0.0001) (**Figure 3I**). This relationship was highly significant for SAT (r=0.87, p=0.0011) (**Figure 3J**) with similar trend for IAT (r=0.49, p=0.11) (**Figure S3K**).

Furthermore, we performed correlation analyses of the ADIPOMAT^lo^ subclusters 6, 8, and 13 with BMI and HbA1c. We observe a significant positive correlation of their abundance with increasing HbA1c levels (**Figure 3K, upper panels**) and a similar trend for correlation with BMI (**Figure 3K, lower panels**).

Together, these findings indicate that the abundance of ADIPOMAT^lo^ adipocytes may be associated with obesity, T2D, and increasing glycemia and may contribute to impaired adipose metabolic function in this setting.

### TSHZ3 is a predicted transcriptional regulator of ADIPOMAT^lo^ mature adipocytes

As differential gene expression and pathway analyses suggested that ADIPOMAT^lo^ cells may represent a metabolically dysfunctional subtype of mature adipocytes associated with metabolic disease, we sought to identify potential transcriptional regulators of this distinct cluster. Predicted regulators of ADIPOMAT^hi^ adipocytes identified by ChEA3 analysis included established key regulators of adipogenesis, such as PPARG, CEBPA, FOXO1, and MEOX1 (**Figure 4A, left**). Interestingly, distinct transcription factors were predicted to mediate gene expression patterns in ADIPOMAT^lo^, including TSHZ3, ZBED6, ZNF469, TWIST2, TCF4, BNC2, TSHZ2, NFIA, FOXD1, and PRRX1 (**Figure 4A, right**); several of these factors are known to be inhibitory for adipogenesis. For example, PRRX1 has been shown to inhibit adipogenesis via activation of the TGFβ signaling pathway^27^, and TWIST2 is known to repress multiple target genes and inhibit adipogenesis^28^. Notably, several predicted regulators of ADIPOMAT^lo^ adipocytes, including TSHZ3 and BNC2, were identified as adipogenesis regulators in our previous pseudotime analysis of *ex vivo* adipose differentiation^29^.

**Figure 4.**
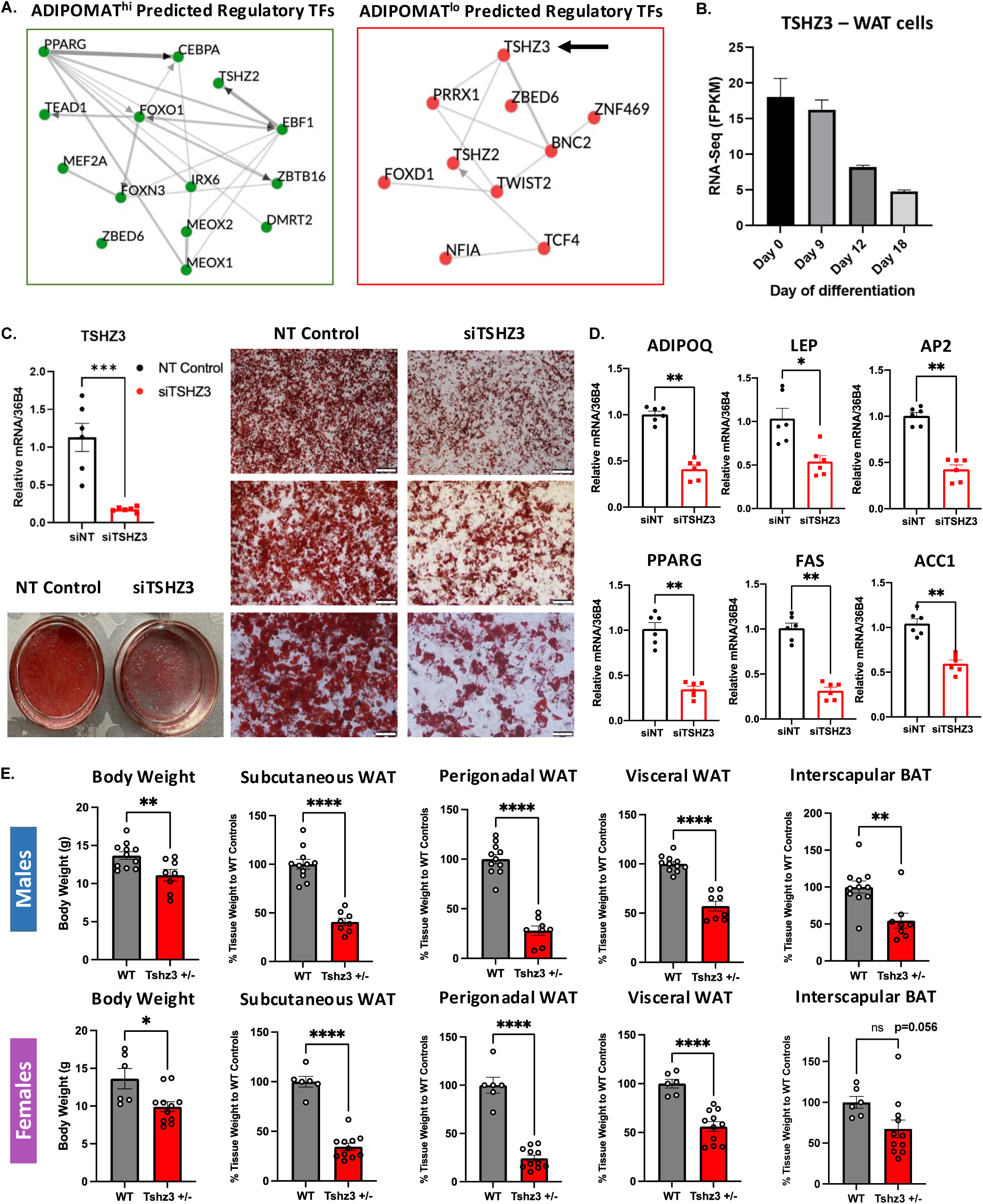
TSHZ3 deletion impairs adipogenesis in human immortalized white adipogenic progenitors and reduces adipose tissue depots in vivo in mice. A. Identification of TSHZ3 as a potential regulator of the transcriptional program of ADIPOMAT^lo^ adipocytes (ChEA3 analysis, input adjusted-p<0.05, log_2_FC>0.75), demonstrating transcription factors predicted to regulate the transcriptional program of ADIPOMAT^hi^ adipocytes (left) or ADIPOMAT^lo^ adipocytes (right). B. TSHZ3 gene expression (normalized FPKM) in human immortalized white adipogenic progenitors and adipocytes at day 0, 9, 12, and 18 of differentiation. C. Efficacy of siRNA-mediated TSHZ3 vs. nontargeted (siNT) knockdown in human immortalized white adipogenic progenitors (RT-PCR, day 0, left), and impact on differentiation (images of Oil Red O-stained cells, right). D. Gene expression of ADIPOQ, LEP, AP2, PPARG, FAS, and ACC1 (RT-qPCR) in human immortalized mature white adipocytes (day 18 of differentiation) in treated with siTSHZ3 or siNT-control on day −2, prior to induction of differentiation. E. Body weight and adipose depot weights in haploinsufficient Tshz3+/lacZ and wild-type control male (upper panels) and female (lower panels) mice. Graphs are presented as mean +/− SEM. * p<0.05, **p<0.01, ***p<0.001, ****p<0.0001.

Given that both TSHZ2 and TSHZ3 were identified as potential transcriptional regulators of the ADIPOMAT^lo^ population, we explored the role of this family in adipogenesis. Previous studies demonstrated siRNA-mediated knockdown of Tshz2 in primary mouse adipogenic progenitors increased expression of Pparg and adiponectin in mature adipocytes, whereas knockdown of the Tshz2-derived circular RNA (circTshz2-1) completely inhibited adipogenesis^30^. Furthermore, recent studies have linked TSHZ3 and insulin resistance in induced pluripotent stem cells (iPSCs)^31^ and two single-nucleotide polymorphisms within the intron of TSHZ3 are associated with T2D^32,33^. Collectively, these data suggest a putative role for TSHZ3 in healthy adipose tissue expansion and its link to insulin resistance.

### TSHZ3 is required for adipogenesis of cultured human white pre-adipocytes

To investigate the role of TSHZ3 in adipogenesis, we first evaluated its expression in adipose-residing cell populations in our snRNA-seq data. TSHZ3 is expressed in adipogenic progenitors (APCs), as well as in mesothelial, endothelial, and immune cells, but not in mature adipocytes (**Figure S4A).** We hypothesized that TSHZ3 regulates early steps of adipogenic differentiation from APC and thus dysregulated function in APC could impair adipogenesis. To test this hypothesis, we performed siRNA-mediated knockdown of TSHZ3 prior to induction of differentiation of human immortalized adipogenic progenitors (day-2) and evaluated full differentiation at days 18-21. These cells robustly express TSHZ3 (**Figure 4B**); in response to experimental TSHZ3 knockdown, expression was reduced by >80% (p<0.001) at day 0 (**Figure 4C**). Lipid accumulation, visualized by Oil Red O staining, was significantly reduced at day 18-21 (**Figure 4C**). In parallel, there was marked reduction (50-60%) in expression of Adiponectin, Leptin, AP2, PPARG, CIDEA, and other key metabolic regulatory genes, confirming reduced adipogenic differentiation with TSHZ3 knockdown (**Figures 4D, S4B**).

Next, we sought to dissect the role of TSHZ3 on adipogenesis in vivo. We generated haploinsufficient *Tshz3* knockout (*Tshz3*^+/lacZ^) male and female mice on a CD1 background and evaluated their adiposity on postnatal days 16, 17, and 18. Strikingly, *Tshz3*^+/lacZ^ male and female mice had significantly lower body weight and dramatically reduced subcutaneous, perigonadal, and visceral white adipose, and interscapular brown adipose tissue depot weights (∼50-60% reduction for each depot), as compared to wild-type littermate controls (**Figure 4E, S4C**). Immunofluorescence of mouse inguinal WAT on embryonic day E18.5 demonstrated that TSHZ3 is primarily co-expressed in perivascular and other adipose-resident EBF2+ cells but minimally co-expressed with PPARG+ cells, suggesting the effects of *Tshz3* deletion may be attributed to its function in EBF2+ adipogenic progenitors (**Figure S4D**). These findings indicate that heterozygous whole-body deletion of *Tshz3* can significantly impair development of adipose tissue - independent of anatomical location, revealing TSHZ3 as a crucial regulator of adipogenesis in vivo. Notably, TSHZ3 expression in adipogenic progenitors (APCs) and mesenchymal-like mesothelial cells (MesoCs2) in another single-nucleus RNA-Seq dataset^34^ of human abdominal visceral (omental) WAT biopsies, was increased in metabolically healthy (MHO) vs. metabolically unhealthy (MUO) - as determined by their insulin sensitivity – in participants with obesity (**Figure S4E**).

To identify early TSHZ3-dependent transcriptional regulation potentially driving its impact on adipogenesis, we evaluated gene expression during early stages of adipogenesis (days 0-8). TSHZ3 expression was low after induction of differentiation (days 2, 5, and 8) in cells treated with the scrambled control siRNA, but was further suppressed in siTSHZ3-treated cells (**Figure 5A**). TSHZ3 knockdown effects on canonical regulators of early adipocyte differentiation^35^ were observed sequentially, with downregulation of CEBPD on day 2, CEBPA and PPARG on days 5 and 8, and of CEBPB on day 8 (**Figure 5A**). Interestingly, ZNF423, which is required for the maintenance of white adipocyte identity^36,37^ was also significantly reduced on day 5 in TSHZ3 knockdown cells (**Figure S5A**). Conversely, expression of NR4A1, a negative regulator of adipogenesis^38^, was significantly increased during late-stage differentiation, potentially as a secondary response to TSHZ3 knockdown (**Figure S4B/S5A**).

**Figure 5.**
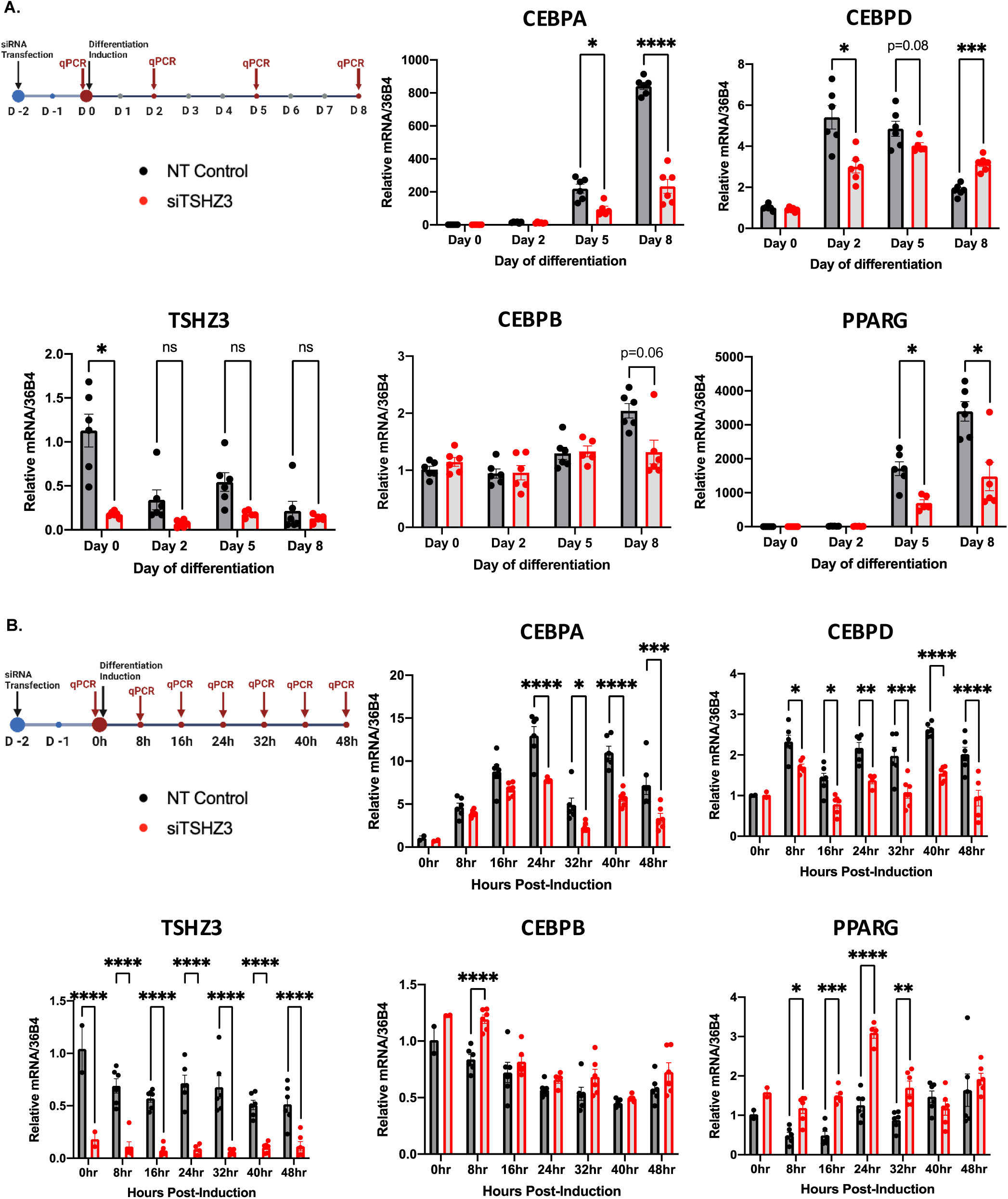
Impact of TSHZ3 KD on differentiation time-course. A. Impact of TSHZ3 knockdown on gene expression of TSHZ3 and canonical adipogenic regulatory genes (TSHZ3, CEBPA, CEBPB, CEBPD, and PPARG) in human immortalized white adipocytes on days 0-5 post-induction of differentiation, B. Impact of TSHZ3 knockdown on gene expression of TSHZ3 and canonical adipogenic regulatory genes (TSHZ3, CEBPA, CEBPB, CEBPD, and PPARG) in human immortalized white adipocytes on early time points (0-48 hours) after induction of differentiation. Graphs are presented as mean +/− SEM. *p<0.05, **p<0.01, ***p<0.001, ****p<0.0001.

To further define the time course of TSHZ3 on very early events in differentiation, we analyzed gene expression at 8-hour intervals for 48 hours after differentiation induction (**Figures 5B, S5A,** heat map in **Figure S5B).** In control cells, we observed the expected coordinated program of adipose differentiation, with progressive and sustained decrease in expression of the inhibitory factors PREF1 and KLF2, and upregulation of CEBPA, CEBPD, EBF1, GJB3, PDGFD, and KLF15. Other early-stage transcription factors such as CEBPB and PPARG family genes did not vary substantially during this time (**Figure 5B and S5B**). By contrast, knockdown of TSHZ3 resulted in complex perturbation of major drivers of the adipogenic program. For example, expression of CREB1, AP1, STAT5A, KROX20, and KLF4 was significantly upregulated at the time of induction (0 hours) (**Figure S5A**). Pro-adipogenic TFs CEBPA, CEBPD, and KLF15 were downregulated at 24 hours (**Figure 5B and S5A/BB**), while CEBPB and PPARG were upregulated (**Figure 5B and S5B**). Additional downregulated transcripts over this early time course included the inhibitory adipogenic transcription factor KLF2 (**Figure S5A-S5B**) and the pro-adipogenic genes EBF1 and KLF15 (**Figure S5A-S5B**).

Given that TSHZ3 KD reduced expression of PPARG, we investigated whether the PPARγ agonist rosiglitazone could rescue adipogenesis in this context. Rosiglitazone was partially effective, as indicated by Oil Red O staining (**Figure S5C**). However, the majority of pro-adipogenic genes (e.g. PPARG, ZFP423) and mature adipocyte markers differentially regulated with TSHZ3 knockdown were not normalized by rosiglitazone, including adiponectin, leptin, and AP2 (**Figure S5D**). Rosiglitazone did significantly attenuate TSHZ3-knockdown-mediated induction of NR4A1 (**Figure S5D**).

Together, these results suggest that TSHZ3 is required for orchestration of the early-stage adipogenic transcriptional program, potentially at the initial clonal expansion phase, with subsequent impact on PPARG-dependent differentiation. Dysregulation of TSHZ3 in adipogenic progenitors dramatically perturbs numerous major TFs that finely tune the initiation of adipogenesis, thus giving rise to adipocytes with impaired mature transcriptomic signature which resemble the ADIPOMAT^lo^ adipocytes.

### Cell-to-cell communication analysis reveals distinct communication profiles from and to ADIPOMAT^lo^ vs. ADIPOMAT^hi^ adipocytes

To identify the unique communication signals among distinct cell types, we employed CellChat^39^ (see Methods), which can deconvolute signals derived from specific cell types (i.e. sources) and targeting others (**Figure 6A and Supplementary Table 3)**. We first assessed communication signals distinguishing the ADIPOMAT^hi^ vs. ADIPOMAT^lo^ subpopulations. Consistent with differential adiponectin expression, ADIPOMAT^lo^ adipocytes provide attenuated adiponectin signaling to endothelial and immune cell populations, as compared to ADIPOMAT^hi^ (**Figure 6B**). Overall, these two cell types had overlapping but distinct patterns of predicted intercellular signaling **(Figure 6C, Figure S6A**). The majority of outgoing signals from ADIPOMAT^lo^ and ADIPOMAT^hi^ were shared (74/88 for ADIPOMAT^hi^ and 74/79 for ADIPOMAT^lo^). However, ADIPOMAT^hi^ adipocytes uniquely provide FGF10, VEGFB, ANGTPL2, CADM1, JAM1, F11R, and JAG1 outgoing signals (**Figures 6D, S6B-S6F**). By contrast, ADIPOMAT^lo^ adipocytes were unique sources of the collagen COL6A3 outgoing signals to other cell types (**Figure 6D, Figures S6B-S6F**). COL6A3 has been previously shown to impair metabolic health of adipose tissue, via restriction of healthy expansion of mature adipocytes^40,41^. Leptin incoming signaling was enhanced in ADIPOMAT^lo^ adipocytes (**Figure 6C, S6A**).

**Figure 6.**
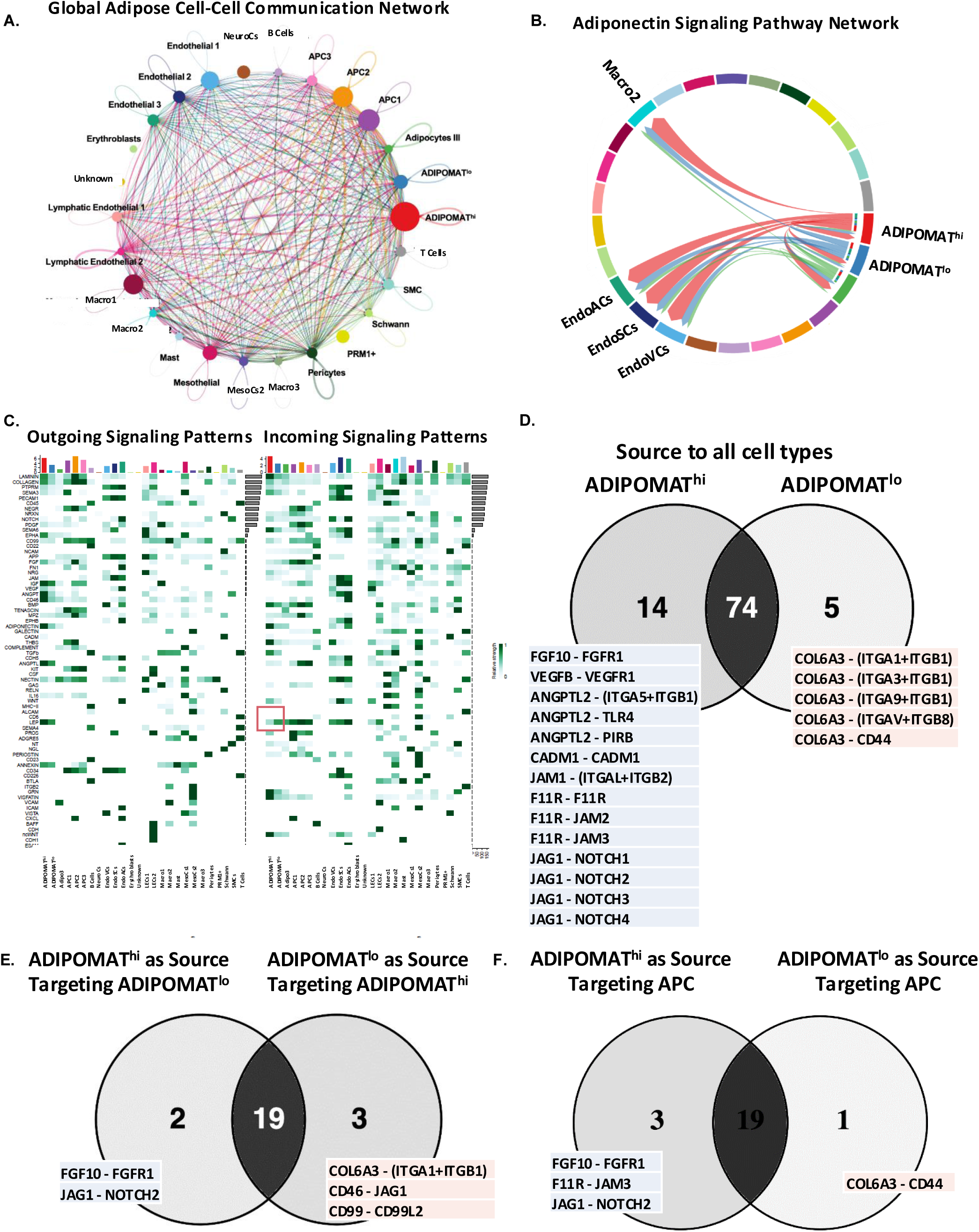
CellChat cell-to-cell communication analysis reveals ADIPOMAT^lo^ adipocytes as sources of COL6A3. A. Global abdominal white adipose tissue cell-cell communication network among white adipose-residing populations. B. Adiponectin signaling pathway network for ADIPOMAT^hi^ and ADIPOMAT^lo^ adipocytes (outgoing) and endothelial and immune cells (incoming). C. Heatmap of outgoing and incoming signaling patterns for white adipose-residing cell populations; red box identifies leptin signaling as incoming signaling pattern increased in ADIPOMAT^lo^ vs. ADIPOMAT^hi^ adipocytes. D. Venn diagram of outgoing signaling patterns from ADIPOMAT^lo^ and/or ADIPOMAT^hi^ adipocytes to all other identified cell types. E. Venn diagram of outgoing signaling patterns from ADIPOMAT^hi^ targeting ADIPOMAT^lo^ and/or ADIPOMAT^lo^ targeting ADIPOMAT^hi^ adipocytes. F. Venn diagram of outgoing signaling patterns from ADIPOMAT^lo^ and/or ADIPOMAT^hi^ adipocytes to adipogenic progenitor cells (APCs).

The majority of signals reflecting crosstalk between mature adipocyte subtypes were shared between ADIPOMAT^lo^ and ADIPOMAT^hi^ adipocytes (**Figure 6E, center**). However, VEGFB, F11R, and JAG1 outgoing signals were uniquely derived from ADIPOMAT^hi^ to target ADIPOMAT^lo^ adipocytes (**Figure 6E, left**), while COL6A3, CD46, and CD99 signals were uniquely derived from ADIPOMAT^lo^, targeting ADIPOMAT^hi^ (**Figure 6E, right**). Communication between mature adipocytes and APCs likely reflects crosstalk crucial for the recruitment of newly differentiated adipocytes from adipogenic progenitors^42^. Signals from ADIPOMAT^hi^ adipocytes to APCs included FGF10, F11R, and JAG1 (**Figure 6F, left**). FGF10 has been previously identified as an important signal stimulating adipogenic progenitors to differentiate into mature adipocytes^43,44^. By contrast, COL6A3 outgoing signaling from ADIPOMAT^lo^ adipocytes targeted APCs (**Figure 6F**). Together these data suggest that increases in collagen products derived from the ADIPOMAT^lo^ adipocyte population and lower pro-adipogenic signals may contribute to both lower differentiation and adverse impact on metabolic health.

## Discussion

Adipose tissue is increasingly recognized as an important metabolic organ essential for healthy metabolism. Disturbances in adipose tissue mass, location, and function yield systemic metabolic disease. Thus, understanding differences in adipose tissue depots, cellular composition, extent of differentiation, and gene expression is essential to dissect its role in the pathogenesis of obesity-associated metabolic disease. While single-cell transcriptomics have been unable to capture the diversity of adipocytes because of their fragile nature, single-nucleus transcriptomics have begun to reveal novel aspects of the molecular heterogeneity of adipose tissue. Our analysis of human adipose tissue obtained from both subcutaneous and intraabdominal depot reveals a subpopulation of adipocytes (ADIPOMAT^lo^) which express lower levels of genes that define maturation and metabolic functionality of adipocytes. Furthermore, this population of cells is more abundant in adipose biopsies from individuals with T2D and is associated with obesity and glycemia-related metabolic phenotypes. Moreover, we identify TSHZ3 as a novel transcriptional regulator, which is expressed in APCs and also modulates adipogenesis in cultured human preadipocytes and modulates adipose mass *in vivo* in mice.

We report a subpopulation of adipocytes with lower expression of multiple transcriptomic markers of mature adipocytes (ADIPOMAT^lo^). ADIPOMAT^lo^ abundance is positively associated with increasing BMI, especially in IAT, suggesting this population may be more prevalent in those with central obesity, who carry greater risk for cardiometabolic disease. This subpopulation is also more prevalent both in individuals with T2D and those with higher glycemia (higher HbA1c). These relationships are not surprising, given that fully differentiated adipocytes orchestrate appropriate physiologic responses to nutrient availability, including both lipogenesis and lipolysis. Moreover, a key secretory product of fully differentiated adipocytes is adiponectin, a systemic adipokine which interacts with cell-surface adiponectin receptors in liver and other key metabolic tissues to regulate systemic glucose and lipid metabolism^45–47^. Our data do not allow us to fully understand the mechanisms mediating the relationship between the ADIPOMAT^lo^ population and metabolic disease; possibilities include reduced secretion of adiponectin per se, altered secretion of other adipokines (e.g. reduced expression of leptin, a key regulator of appetite and lipolytic signaling), or impaired lipogenic capacity – as evident by significant downregulation of the lipogenesis-associated genes ACACB and GPAT3 - in cells with incomplete adipogenesis or de-differentiation. In turn, reductions in lipogenic capacity would be expected to limit lipid storage in adipose tissue, thus increasing lipid overflow to non-adipose tissues such as liver and muscle, promoting systemic insulin resistance and disease risk^11^.

One potential insight into additional mechanisms by which ADIPOMAT^lo^ adipocytes may regulate systemic metabolism comes from the cell-cell communication analyses. While this method may be somewhat susceptible to background noise^48^, we find that ADIPOMAT^lo^ - but not ADIPOMAT^hi^ - adipocytes are a source of collagen and other profibrotic and proinflammatory signals which may signal in a paracrine manner to APC and other adipose-resident cell types, limiting adipogenesis. Indeed, prior studies have demonstrated that COL6A3 restricts healthy expansion of mature adipocytes in response to positive energy balance^40,49^. Such a “barrier” of healthy expansion confers both mechanical stress and hypoxia to mature adipocytes, promoting a pro-inflammatory phenotype also linked to metabolic disease^49^. In addition, ADIPOMAT^lo^ adipocytes (in contrast to ADIPOMAT^hi^) - are not a source of FGF10 signaling to APC and other cell type targets. This is an exciting finding as prior studies have shown that FGF10 signaling is crucial for recruitment of APC and their differentiation into mature adipocytes^50^. Taken together, these data suggest that a higher abundance of ADIPOMAT^lo^ adipocytes may not only promote proinflammatory signaling but also reduce recruitment of adipogenic precursors, further limiting their differentiation into metabolically active, healthy adipocytes. Such dysfunctional adipocyte expansion in response to ADIPOMAT^lo^-mediated signaling ultimately may ultimately contribute to development of metabolic disease.

Another significant finding of our study is the identification of the transcription factor TSHZ3 as a key regulator of human adipogenesis. While multiple transcription factors are putative upstream regulators of the ADIPOMAT^hi^ vs. ADIPOMAT^lo^ transcriptomic signatures, TSHZ3 was unique to the ADIPOMAT^lo^ population. Given that TSHZ3 expression is higher in APCs that are more likely to give rise to ADIPOMAT^hi^ adipocytes and lower in progenitors that are more likely to give rise to ADIPOMAT^lo^ adipocytes, we hypothesized that (1) TSHZ3 is crucial for adipogenesis, and (2) reduced TSHZ3 expression in APC precursors of ADIPOMAT^lo^ adipocytes could impair adipogenic potential. Moreover, two intronic SNPs have been linked to T2D^32,33^. Our data demonstrate that experimental knockdown of TSHZ3 had a striking effect on adipogenesis in cultured human preadipocytes, reducing the formation of adipocytes and adipocyte-specific gene expression by more than 50%. These effects were most prominent in early stages of differentiation, even prior to induction of the canonical early adipogenic transcription factors CEBPB and CEBPD. Differences between early (within 48h post-induction) and later (day 5) expression patterns of PPARG may be related to direct effects of TSHZ3 to repress PPARG (a predicted direct target gene of TSHZ3 based on binding motif analysis using the TF2DNA database) and/or indirect effects, mediated by reduced expression of CEBPs during early adipogenesis^51^. Our in vivo studies further support the role of TSHZ3 as a crucial regulator of adipogenesis, as seen in the dramatic reduction in adipose tissue mass in mice with heterozygous whole-body constitutive deletion of the *Tshz3* gene.

We observe a sub-population of beige thermogenic cells present in both subcutaneous (∼3% of ADIPOMAT^hi^ mature adipocytes) and intraabdominal (∼9-10% of ADIPOMAT^hi^ mature adipocytes) depots. These cells express established transcriptional regulators of the thermogenic program (e.g. EBF2, PPARGC1A, and PRDM16)^52–54^ as well as newly identified marker transcripts such as DPP6, ZNF804A, GIPR, and KCNJ3. While recruitment of beige adipocytes appears to be limited to the subcutaneous depot in mice^55,56^, these intriguing data indicate that human visceral adipose also has the capacity to acquire thermogenic properties with impact on systemic metabolism. In agreement with our data, “beiging” of abdominal white adipose tissue has recently been described in human visceral adipose tissue^15^. Future studies will be required to determine whether transcripts expressed in these cell populations can serve as markers or novel targets for regulating thermogenesis and systemic metabolism.

We acknowledge several limitations of our study. First, the overall number of samples (22) was relatively limited. Donors differ according to multiple factors, including genetic sequence; between-donor differences could limit our ability to identify additional transcriptomic phenotypes distinguishing adipose-resident cell populations and subpopulations. However, a robust number of nuclei was captured from each tissue sample, potentially mitigating the impact of sample variance, thus allowing us to identify new subpopulations of adipocytes. Secondly, the number of participants with T2D was low; it will be necessary to validate associations between ADIPOMAT^lo^ cell populations and glycaemia in a larger cohort. Third, the majority of biopsy donors had high BMI and obesity; further studies will be required to assess abundance of ADIPOMAT^lo^ adipocytes in individuals across a broader range of adiposity and metabolic health. Fourth, a challenge of the field is to make a distinction between the identification of a cell subtype vs. a cell state. Since our findings are based on a “snapshot” of human adipose biology and do not include biopsies from various timepoints, we cannot conclusively state whether the identified ADIPOMAT^hi^ and ADIPOMAT^lo^ adipocytes represent distinct cell subtypes or cell states. Whether ADIPOMAT^lo^ adipocytes represent an intermediate stage of adipocyte differentiation/adipogenesis in progress, cells with intrinsic impairments in adipogenesis which limit differentiation, or mature adipocytes which have been partially de-differentiated will be an important question for future studies investigating time-dependent responses of cell populations in response to stimuli such as diet. Finally, we acknowledge that despite data from both in vitro siRNA and in vivo deletion of TSHZ3 supporting an important role for this transcription factor in adipogenesis, proof of specificity of TSHZ3 within the ADIPOMAT^lo^ population will require analysis of adipocyte populations in lineage tracing models and in response to tissue-specific and temporal disruption of TSHZ3 transcriptional activity, studies beyond the scope of the current manuscript.

In summary, our analysis of the nuclear transcriptome in human IAT and SAT represents an important resource which can contribute toward a deeper understanding of human adipose-residing cells, adipocyte biology, and depot-specific regulation of the complex processes that govern adipogenesis in vivo. Furthermore, we identify novel metabolically distinct subtypes of adipocytes based on low vs. high expression of genes that mark adipocyte maturity, demonstrate differential abundance of this population in relation to BMI and glycaemia, and identify TSHZ3 as a novel upstream regulator of adipogenic differentiation in APC which may contribute to subpopulation differences in mature adipocyte function and metabolic regulation (**Figure 7**).

**Figure 7.**
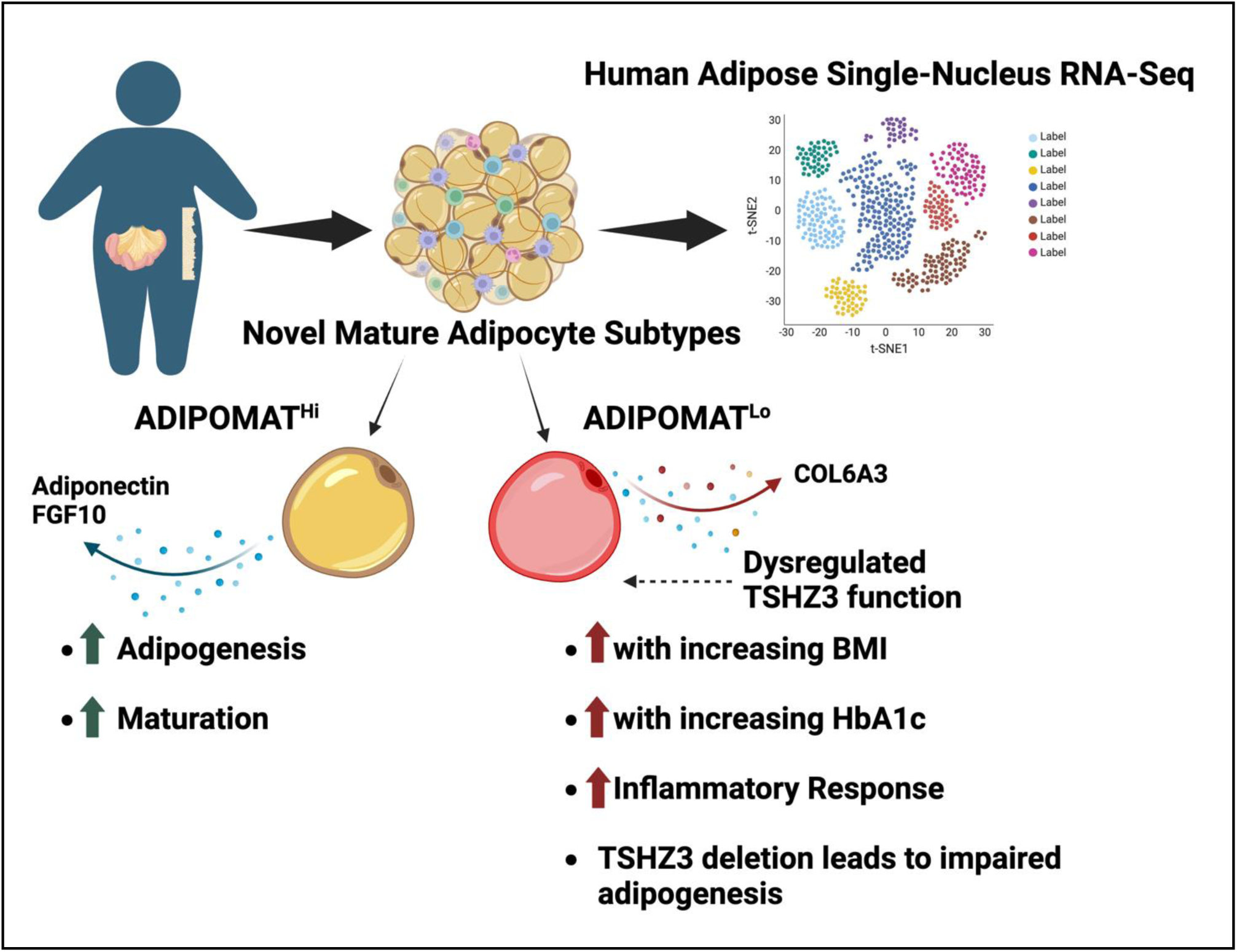
Summary figure. Comprehensive human cell atlas of subcutaneous and intraabdominal white adipose tissue depots, capturing canonical cell types and novel subpopulations. We identify two major mature adipocyte subtypes, characterized by high and low expression of genes that mark maturation level of adipocytes (ADIPOMAT^hi^ and ADIPOMAT^lo^). ADIPOMAT^lo^ adipocytes have reduced adipogenic and increased pro-inflammatory and pro-fibrotic transcriptomic signatures, and their abundance is positively associated with BMI and HbA1c. We identify TSHZ3 as a key transcriptional regulator of early steps of human adipogenic progenitor differentiation which may contribute to the ADIPOMAT^lo^ transcriptome.

## Methods

### Collection of human white adipose tissue biopsies

White adipose tissue biopsies were collected from two locations (intraabdominal or IAT and subcutaneous abdominal or SAT) during laparoscopic elective surgery at Brigham and Women’s Hospital (BWH, Boston, MA). For the sleeve gastrectomy, LigaSure bipolar cautery was used and for the gallbladder surgical procedure, hook cautery was used for tissue extraction and biopsy acquisition. Subcutaneous adipose tissue biopsies were obtained from the abdominal area, whereas intraabdominal adipose tissue biopsies were obtained from the gastrocolic or omental anatomical area (**Supplementary Table 4**). Tissues were snap-frozen in dry ice and kept frozen in −80° C until further processing. The study was approved by the Institutional Review Board at Brigham and Women’s Hospital, and all donors provided written informed consent.

### Single-Nucleus Isolation, Sample Preparation, RNA-Sequencing, Analysis (Batch Correction, Integration, Clustering) and Visualization of Human WAT Biopsies

Intact white adipose tissue-derived nuclei were isolated using a glass-on-glass homogenizer protocol. In detail, ∼100mg of WAT were placed on a petri dish with tissue lysis buffer (Tris-HCl (pH 7.4, 10mM), MgCl_2_ (3mM), NaCl (10mM), NP40 (0.1%), and RNase inhibitors (Takara, Recombinant RNase Inhibitors, 1U/uL)) until thawed. Thawed samples were placed on cold-block and cut with sharp scissors for 3 minutes. Samples were subsequently transferred to a glass homogenizer in a total volume of 7 mL of lysis buffer. Samples were homogenized ∼20 times (10 times with “loose” pestle, 10 times with “tight” pestle) and incubated on ice for 6 minutes. After incubation, 5 mL of “blocking buffer” (1% BSA in PBS + RNase inhibitors (Takara, Recombinant RNase Inhibitors, 1U/uL)) was added and samples were centrifuged at 500g for 5 minutes at 4°C. After centrifugation, supernatant was aspirated, and pellet was washed in blocking buffer and transferred to a new Falcon tube. After a total of 2 washes with blocking buffer, nuclei were stained with Trypan Blue and counted and an average of 16,000 nuclei were loaded onto each lane of a 10x Chromium chip platform, per manufacturer’s instructions.

10x Genomics libraries were sequenced at a mean depth of 50,000 reads per nucleus (Illumina NovaSeq, BioHub, San Francisco). Data were filtered using CellRanger^57^ and CellBender^58^ to remove low-quality nuclei. In addition, nuclei and clusters that had very low number of unique markers and/or high number of mitochondrial (>5%) and/or ribosomal markers were removed, as these metrics are associated with poor nuclei transcriptome quality. Background removal was performed using SoupX^59^. DoubletFinder was applied to remove doublets.

Batch effects from different flow cells and different chips are likely to be minimal (Tech Note: https://kb.10xgenomics.com/hc/en-us/articles/115003122252). However, due to experimental constraints related to access to human biopsies and lengthy nuclear isolation protocols, nuclei from adipose biopsies from different donors were isolated and processed on the 10X Chromium platform for subsequent library preparation on different days. Likewise, libraries were submitted for Illumina Sequencing on different days. Hence, for downstream data integration, each sample was defined as a batch.

For each biopsy sample, CellRanger^60^ was used to process sequencing and generate raw (raw_feature_bc_matrix.h5) and filtered (filtered_feature_bc_matrix.h5) count matrices. The “chemistry” parameter for running CellRanger was set as “Single Cell 3’ v3”. The resulting raw count matrix (raw_feature_bc_matrix.h5) was then input into CellBender (v.2)^58^ for background ambient RNA removal with two output files that would be further used in the later preprocessing steps. When running CellBender, “expected-cells” parameter was set to be the number of barcodes in filtered_feature_bc_matrix.h5 file, and “total-droplets-included” parameter was set to be 3 times of the expected-cells number. The two output files are: 1. full count matrix (output.h5) that contains all the original droplet barcodes, and 2. filtered count matrix (output_filtered.h5) with barcodes which were determined by CellBender to contain cells. The output_filtered.h5 was then used for quality control. Doublet detection and filtering was performed using scds^61^, and low-quality cells were identified based on mitochondrial transcripts percentage level, using 1% as a threshold. We used default parameters running scds. Barcodes that were flagged as either doublets or low quality were then filtered out from output_filtered.h5. As an additional quality filter, barcodes that did not appear in filtered_feature_bc_matrix.h5 were also removed in output_filtered.h5. SoupX^59^ was then used to estimate and remove cross contamination of the transcripts across cells. A list of broad cell type marker genes and barcodes further filtered output_filtered.h5 were the input for SoupX and output soupx_adjusted_counts.h5ad.

After background removal with SoupX, each sample has a corrected expression count matrix output soupx_adjusted_counts.h5ad. Samples were then integrated and batch corrected with SCVI^62^ which produces a low dimensional latent representation of the gene expression matrix. A first-round cell clustering was performed on SCVI’s low dimensional representation. The resulting integrated data across samples are available on CellxGene [https://cellxgene.cziscience.com/collections/6b701826-37bb-4356-9792-ff41fc4c3161]. The pre-processing process is shown in Supplementary Figure 1A.

All the parameters chosen for above tools are specified in the scripts provided. CellRanger parameter specification is in run_cellranger_multiplesample.sh file. CellBender parameter specification is in pipeline_beta.sh file. scds parameter specification is in QC_scds.R file. SoupX parameter specification is in QC_soupx.R file, and the marker gene list as input for SoupX is in Arionas_marker_gene_list_noquotes.csv file. SCVI parameter specification is in integrate_soupx_output.py file.

scVI was used for integration of high-quality nuclei from all 22 biopsy samples obtained from different donors, depots, and experimental days^16^. In scVI-tools, variable genes from each batch are identified and their intersection is used for integration. Utilization of genes that are commonly variable across batches helps remove batch-specific variation due to batch-specific gene expression. To assess whether integration using scVI-tools worked reasonably well, we referred to the latent space inferred for each biopsy (**Figure S1B**); this analysis demonstrated that all 22 biopsies were relatively mixed in latent space. The integration was assessed based on the calculation of LISI scores across tissue and samples. We estimated average LISI scores using the R package LISI v1.0^63^ by adding embeddings of the UMAPs along the metadata.

Integrated data were clustered and visualized using unsupervised approaches in Seurat^64^ (Version 4.9.9.9049) in R (version 4.3.1). Individual gene expression was plotted using the FeaturePlot or DotPlot functions in Seurat. Initially, 27 clusters were identified, using 1:11 dimensions and 0.9 resolution and UMAP plot was visualized using default parameters. However, further analysis identified cluster 3 as a low-quality population, due to expression of markers expressed in multiple cell types (e.g. adipocytes, endothelial cells, and immune cells), and was thus removed from further analysis. FindAllMarkers function (with default parameters using log10k-normalized data) was used to identify markers for each cluster. For the reference mapping approach, anchor cells were identified between the reference Emont et al. dataset^15^ and our snRNA-seq dataset in a shared low-dimensional space using overlapping highly variable features. These anchors were used to project our snRNA-seq dataset onto the reference, transferring cell type labels and calculating prediction scores using Seurat v4.4. Prediction scores and top cluster markers were used together for cell type annotations.

Differential gene expression analyses (using the FindMarkers function with default parameters on log10k-normalized data), identification of unsupervised cluster markers, and visualization of violin plots, dot plots, feature plots, and UMAPs were performed using Seurat (Version 4.9.9.9049) in R (version 4.3.1).

ADIPOMAT^lo^ signature was created separately for IAT and SAT based on the upregulated genes shown in Figure S2L. The depot specific signature was separately used to create module scores for VAT and SAT dataset from Lazarescu et al^25^.

### Pathway Enrichment Analysis

Pathway enrichment analyses were performed using ENRICHR (https://maayanlab.cloud/Enrichr/)^65^ and MsigDB^23^. Genes that were identified via differential gene expression analysis or via unsupervised cluster marker analysis and had log_2_FC > 0.5 and adjusted p< 0.05 were used as input.

### Predicted Regulatory Transcription Factor Analysis

Predicted regulatory transcription factor analysis was performed using ChEA3 (https://maayanlab.cloud/chea3/)^66^. Genes that were identified via differential gene expression analysis or via unsupervised cluster marker analysis and had log_2_FC > 0.75 and adjusted p-value < 0.05 were used as input.

### VISION analysis

The VISION tool was applied to calculate an adipogenesis signature score (based on the “Hallmark Adipogenesis” MsigDB^23^) for each cluster in the integrated Seurat object, as described in DeTomaso et al^22^.

### Cell-Cell Communication (CellChat) Analysis

Cell-cell communication (i.e. ligand-receptor interaction) analysis was performed using the R package CellChat^39^. Using normalized gene expression and cell clusters derived from the Seurat object, we computed the communication probability and inferred cellular communication network among the cell clusters. Data are reported in tables (**Supplementary Table 3**) and include ligand-receptor interaction and the summarized pathway level statistics; the column “prob” represents the communication probability (i.e. strength of interaction) and the column “pval” represents the p-value level of significance.

The aggregated cell-cell communication network is visualized via circle plots, which show ligand-receptor interactions weights of the interaction from each individual cluster (the signal sender) to other clusters (the signal receivers).

To visualize the communication network at a single pathway level, we selected the ADIPONECTIN pathway due to its importance in adipocyte function. The contribution of each ligand-receptor pair is visualized in circle plots (in which mature adipocyte clusters are the signal senders). Analysis at the system level for signaling roles i.e. sender, receiver, mediator, and influencer for each pathway is shown in a heatmap demonstrating the relative strength of each signaling pathway in terms of outgoing and incoming patterns. To further dissect the patterns of outgoing and incoming signals, we used the “select-K” function to infer the number of patterns K. We show two measures to select K (**Supplementary Table 3**). We choose K as the number of patterns immediately before the sudden drop of the two measures (Cophenetic and Silhouette). The heatmaps of outgoing and incoming signaling patterns show the contribution of each cluster (cell patterns) and each signaling pathway (communication patterns).

### In vitro mature human adipocyte differentiation

Human immortalized mature adipocytes were differentiated in vitro from adipogenic progenitors derived from the subcutaneous neck region^67^. Adipogenic progenitors were cultured in high-glucose (4.5g/L) DMEM medium supplemented with 10% fetal bovine serum (FBS) and 1% penicillin/streptomycin (growth medium). All lines were maintained below confluence by splitting every 2-3 days. To induce differentiation, growth medium was replaced with induction medium, containing human insulin (0.5uM), dexamethasone (0.1uM), IBMX (500uM), indomethacin (30uM), pantothenate (17uM), and biotin (33uM) added to the growth medium, upon confluence (defined as day 0 of differentiation). Medium was refreshed every 72 hours, until day 18 where the maximum percentage of adipocyte differentiation is achieved. At the end of the experiment, differentiation medium was removed, cells were washed with PBS, and subsequently frozen at −80°C (for RT-qPCR analysis) or fixed with 4% formaldehyde (for Oil Red-O staining), until further processing and analysis.

### siRNA-mediated knockdown in vitro in human adipocytes

Human immortalized adipogenic progenitors (100,000 cells/well) were transiently transfected with 10nM of siRNAs targeting human TSHZ3 (Dharmacon IDT) using the transfection reagent Dharmafect (15uL of Dharmafect per 1mL of OptiMEM) in OptiMEM (15uL of Dharmafect per 1mL of OptiMEM). Cells were incubated with transfection mix and growth medium for 48-60 hours prior to induction of differentiation.

### RNA extraction – Reverse Transcription

Cells were lysed in Trizol and the aqueous phase was collected after chloroform addition. RNA was pelleted using isopropanol, treated with Dnase I (NEB) and precipitated using sodium acetate and ethanol. Subsequently, reverse transcription was performed to generate cDNA (High-Capacity cDNA Reverse Transcription Kit, Applied Biosystems).

### qPCR gene expression analysis

cDNA was diluted to 5ng/uL and quantitative PCR (qPCR) was performed in a QuantStudio 6 real-time PCR system using Fast SYBR Green qPCR Mastermix (Applied Biosystems) with validated primers. Gene expression analysis utilized QuantStudio software (Applied Biosystems); relative mRNA concentrations normalized to the expression of the housekeeping gene were calculated using the ΔΔCt method.

### Mouse strains

The *Tshz3^lacZ^* mouse line was generated on a CD1 background as previously described^68^. Wild type (WT) and *Tshz3^+/lacZ^* male and female mice were obtained by crossing heterozygous *Tshz3^+/lacZ^* male mice to WT females; WT littermates served as controls. Animals were maintained on a 12-hour/12-hour light/dark cycle in a temperature and humidity-controlled environment with food and water *ad libitum*. Experimental procedures were in agreement with the recommendations of the European Communities Council Directive (2010/63/EU).

### Mouse tissue preparation

WT and *Tshz3^+/lacZ^* mice at postnatal day 16-18 were euthanized by cervical dislocation. White and brown adipose tissues (WAT and BAT) were dissected and immediately weighed on an analytical balance (AE260 Delta Range, Mettler Toledo, Greifensee, Switzerland).

### Oil Red O staining

Oil-Red O was used to visualize and quantify lipid droplets and lipid content. After removing the supernatant, cells were fixed with 4% formaldehyde solution in PBS for 30 minutes at room temperature. Oil-Red O (stock solution of 0.5g Oil Red O powder dissolved in 100mL isopropanol) was diluted 1:4 in water (working stock). Cells were incubated with the working stock for at least 1 hour at room temperature, followed by multiple washes with PBS. Images were acquired using a conventional light microscope. Lipids were extracted using isopropanol and absorbance was quantified using QuantStudio spectrophotometer at 518 nm.

### Statistics and Graphs

For in vitro and in vivo experiments, GraphPad Prism (version 10.1.2) was used to generate graphs and perform statistical analyses. Unpaired two-tailed t-test was performed for two-group comparisons, and one-way ANOVA for comparisons of three or more groups, using Tukey’s test for multiple-comparison corrections. Simple linear regression analyses (Pearson r and p-value for correlation are calculated) were also performed using GraphPad Prism (version 10.1.2).

## Data Availability

Single-nuclei RNA-Sequencing datasets generated and used in this study will be deposited into dbGaP upon acceptance. All data that support the findings of this study are included in this paper within supplementary information files. Source data are provided with this paper. Data and visualization are also available via the CellxGene platform: https://cellxgene.cziscience.com/collections/6b701826-37bb-4356-9792-ff41fc4c3161.

## Code Availability

CellRanger, CellBender, scVI and Seurat analyses were performed following the standard tutorials of the respective Python and R packages. Python and R code used in this study will be available via GitHub (https://github.com/jdreyf/efthymiou-patti-adipose-snRNAseq) upon acceptance.

## Acknowledgements

We thank the Chan-Zuckerberg Initiative (CZI) and the Swiss National Foundation (SNSF) for funding. We would like to thank the BioHub (San Francisco, CA) and the Harvard University Bauer Core (Cambridge, MA) for RNA-sequencing, and specifically thank Norma Neff and the CZ Biohub Sequencing Team for lending support with sequencing snRNA-seq libraries. A.S. is a Chan Zuckerberg Biohub Investigator. Authors acknowledge the Howard Hughes Medical Institute for funding A.S.B. through the Hanna H. Gray Fellows program. We also thank Dr. Jonathan Dreyfuss and Dr. Hui Pan (Joslin Bioinformatics Core) for CellChat analyses We would also like to acknowledge support from the Joslin Diabetes Center Diabetes Research Center, NIH P30 DK036836.

## Author Contributions Statement

V.E. and M.E.P. designed the study; V.E., Y.H.T., A.S., and M.E.P. supervised the experiments; V.E., S.D.K., and F.S. performed the single-nuclei RNA-Seq 10x experiments; V.E., W.A., and L.P. performed in vitro human adipocyte experiments; S.Y. preprocessed all FASTQ files and generated count matrices; S.Y., R.G., and A.S.B. performed initial quality control; S.Y., and R.G. performed initial matrix processing and cell type clustering; Ad.Gh. and V.E. performed follow-up clustering and cell type assignment; Ad.Gh., V.E., S.D.K, An.Gu., and H.C. analyzed differences between the two adipocyte subclusters and performed VISION analyses; Y.B., X.C., and L.F. performed in vivo experiments in *Tshz3*^+/lacZ^ mice. Y.H.T. and A.S. provided resources; Y.H.T. provided immortalized human adipogenic progenitor cell lines; A.V. provided human abdominal adipose tissue biopsies; V.E. and M.E.P. wrote the manuscript; all authors reviewed and edited the manuscript.

## Inclusion and Ethics Statement

The paper adheres to the Nature Portfolio journals’ authorship policy.

## Supplementary Figure Legends

**Supplementary Figure 1.**
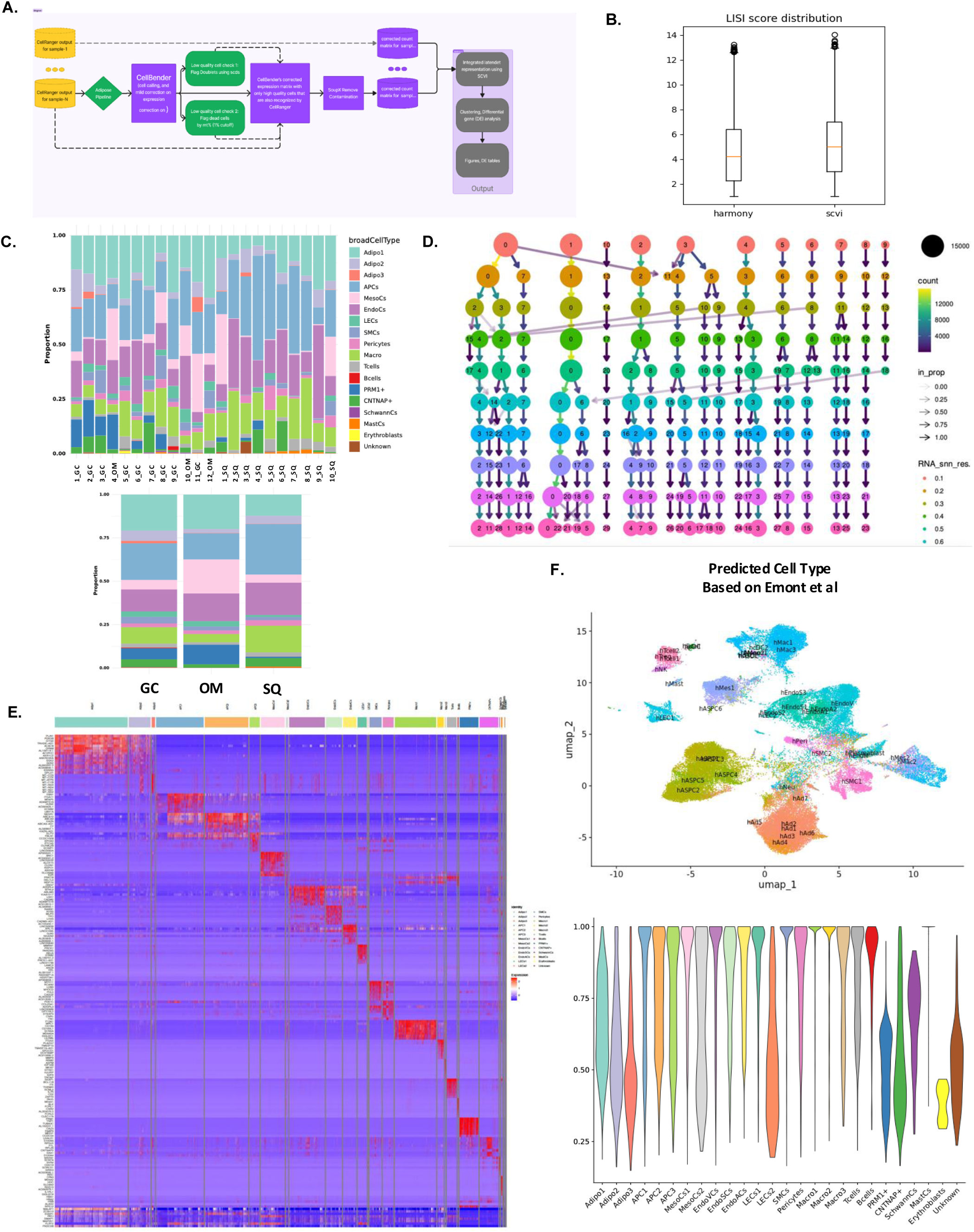
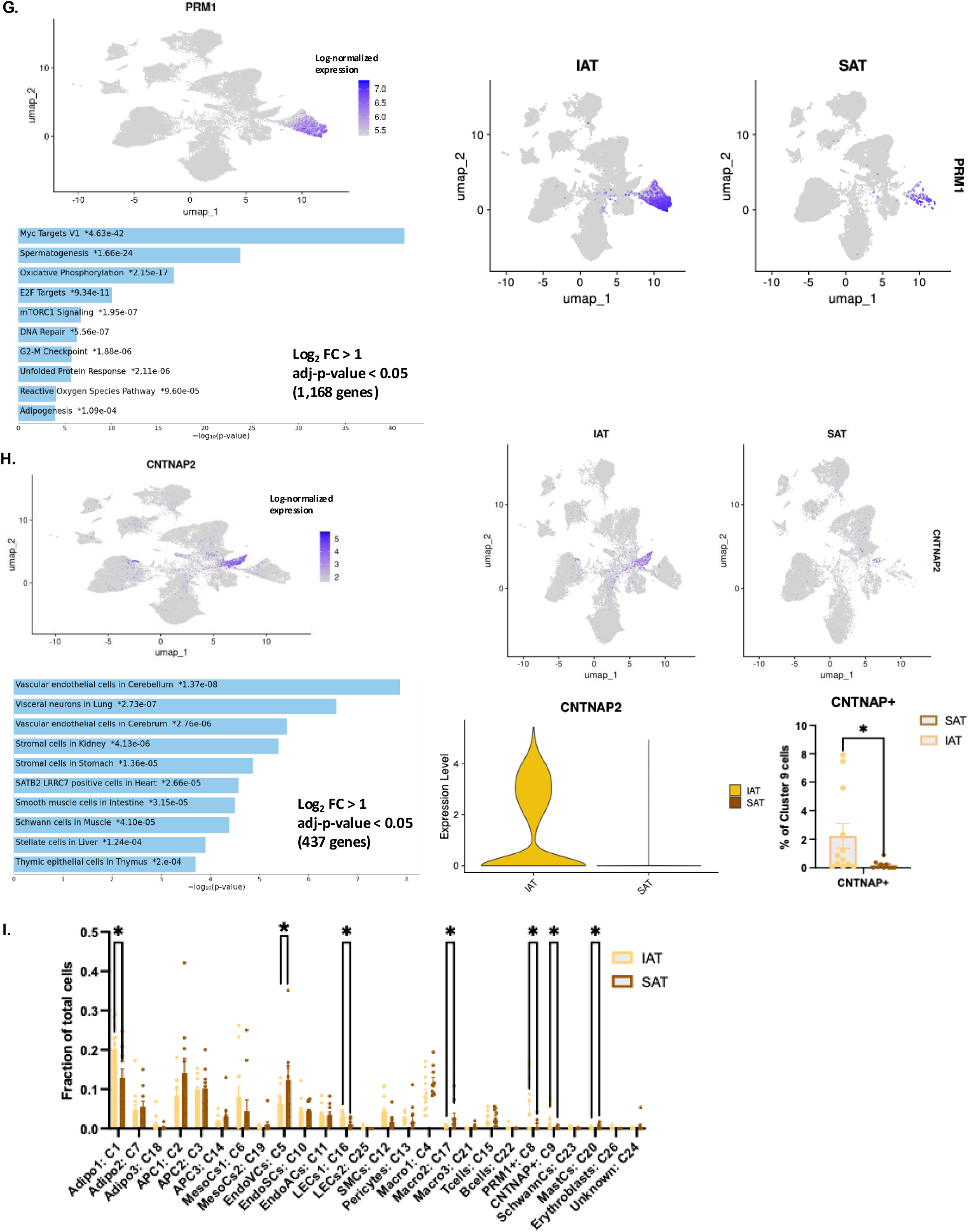
A. Flow chart of analysis pipeline. B. LISI score distribution, calculated using Harmony (left) or scVI (right). C. Stacked BarPlot showing all identified broad cell types, divided by individual donor (upper). Stacked BarPlot showing all identified broad cell types, divided by anatomical depot (gastrocolic AT (GC), omental AT (OM) or subcutaneous AT (SQ) (lower). D. Clustering tree analysis. E. Heatmap showing top markers of all identified cell clusters, F. UMAP plot showing predicted cell types of all cell clusters, annotated based on reference mapping to the Emont et al dataset^15^ (upper) and cell type prediction score of reference mapping per cell cluster (lower), G. UMAP plot showing expression of PRM1 for all biopsies (upper left panel), expression of PRM1 divided by anatomical location (IAT: left, SAT: right) and pathway enrichment ontology analysis of PRM1+ cell population (lower). Blue color of the bar signifies significant enrichment for the respective pathway and length of the bar signifies −log_10_ p-value, H. UMAP Plot showing expression of CNTNAP2 for all biopsies (upper left panel), expression of CNTNAP2 divided by anatomical location (IAT: left, SAT: right), and pathway enrichment ontology analysis of NeuroCs cell population. Blue color of the bar signifies significant enrichment for the respective pathway and length of the bar signifies −log_10_ p-value (lower left panel). Violin Plot showing expression of CNTNAP2 divided by anatomical location (IAT: left, SAT: right) and percentage of CNTNAP+ cells within NeuroCs cell population (lower right panels). I. Percentage of all cell clusters divided by anatomical location (IAT: left, SAT: right). Graphs are presented as mean +/− SEM. *p<0.05.

**Supplementary Figure 2.**
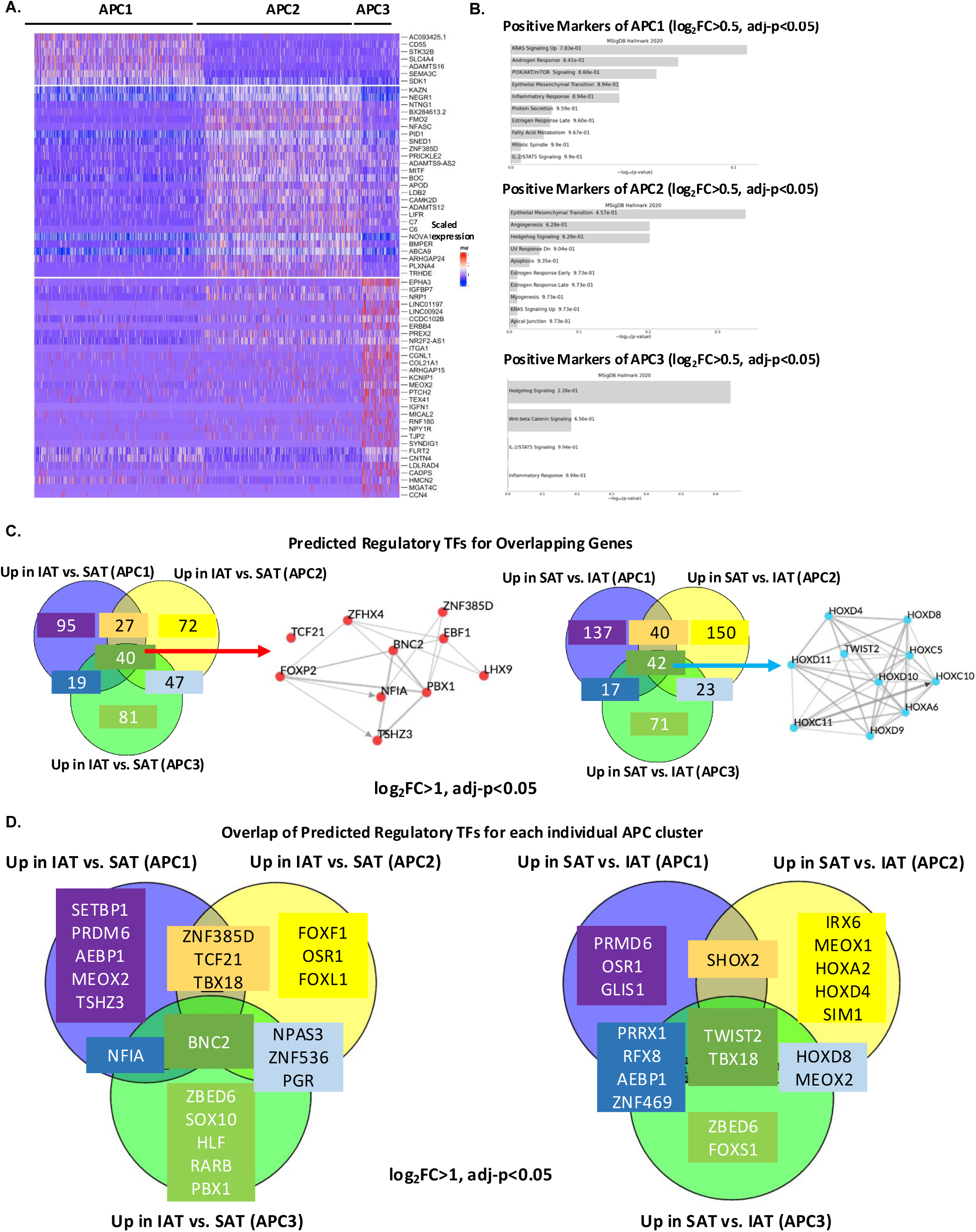
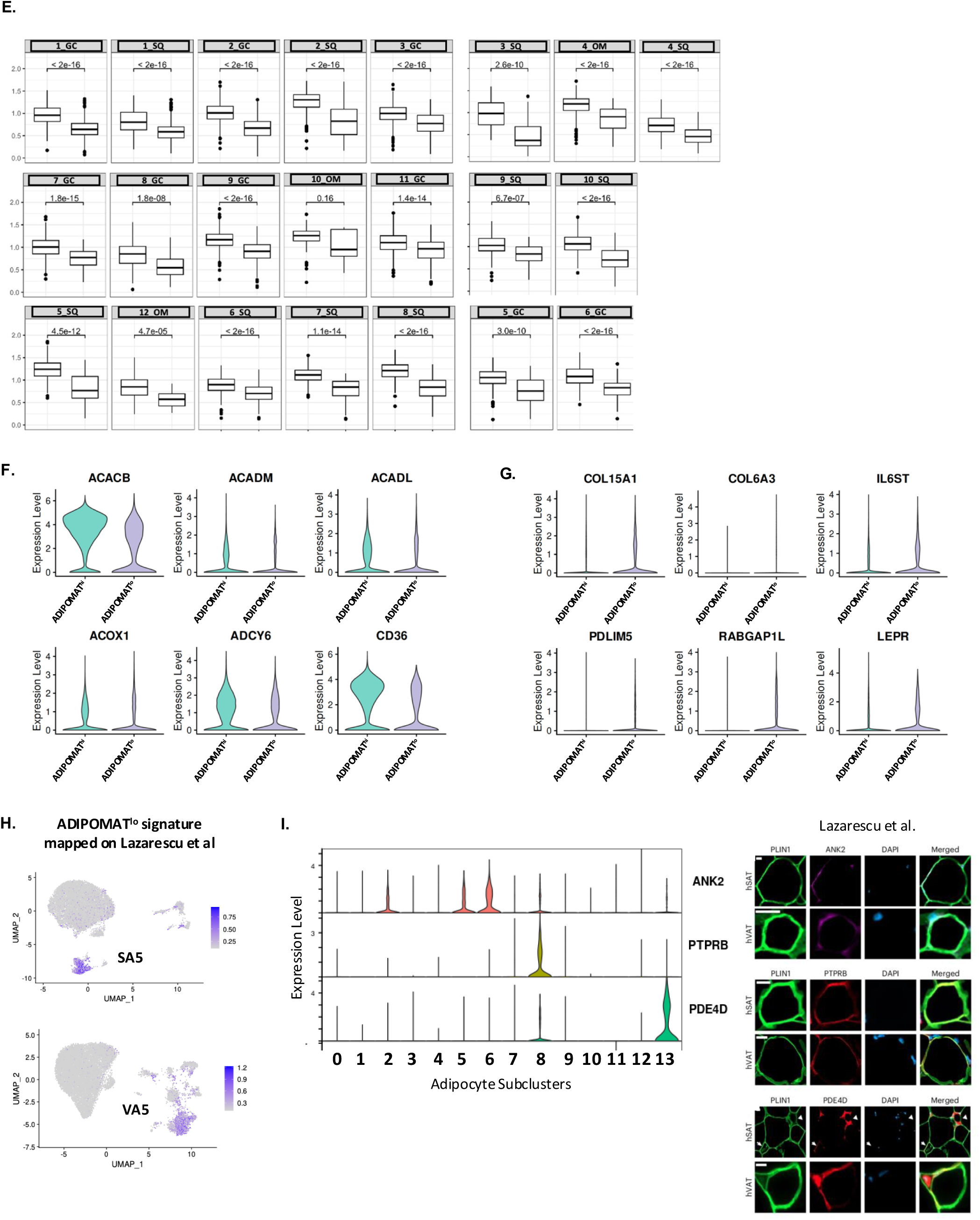
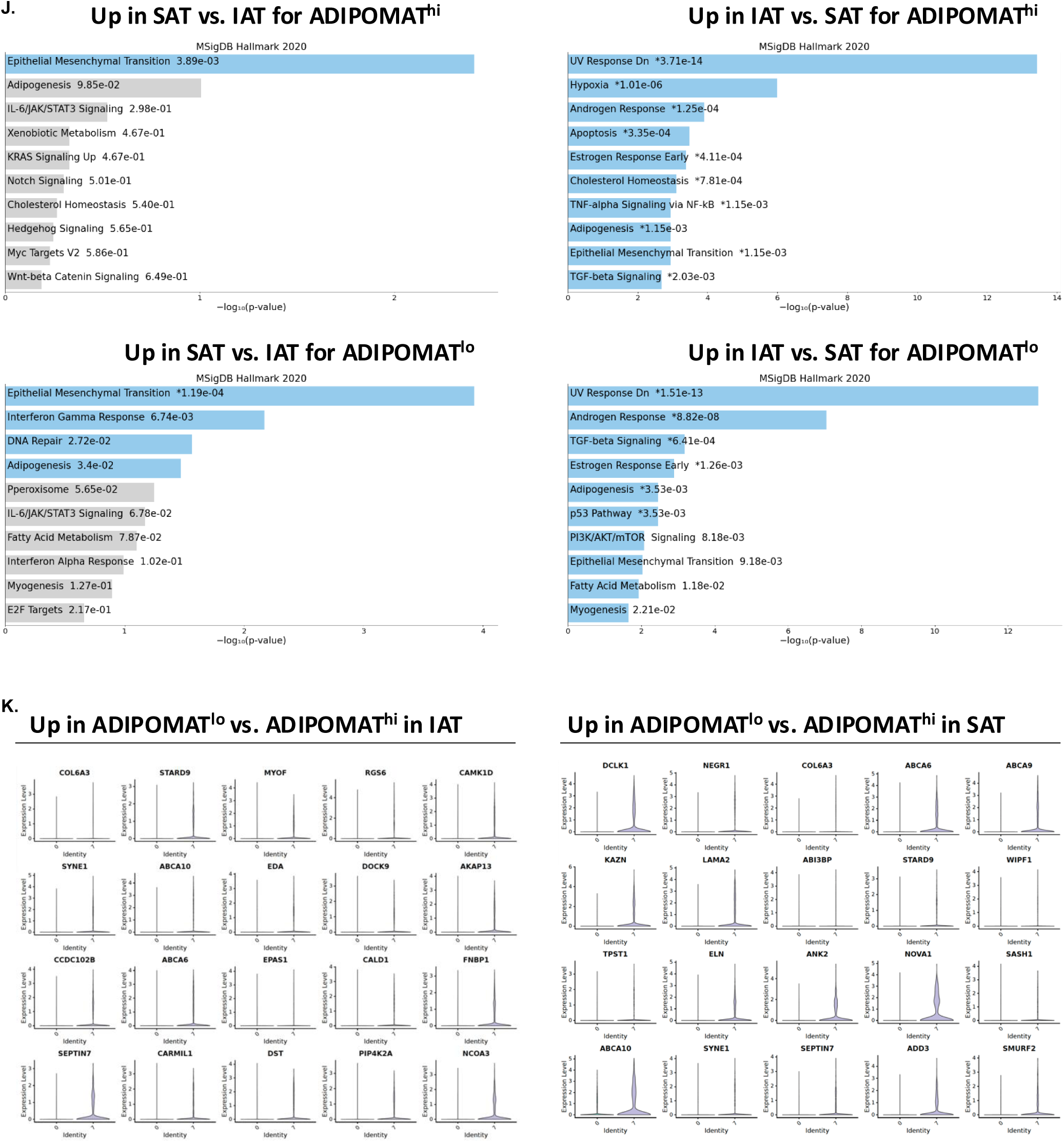
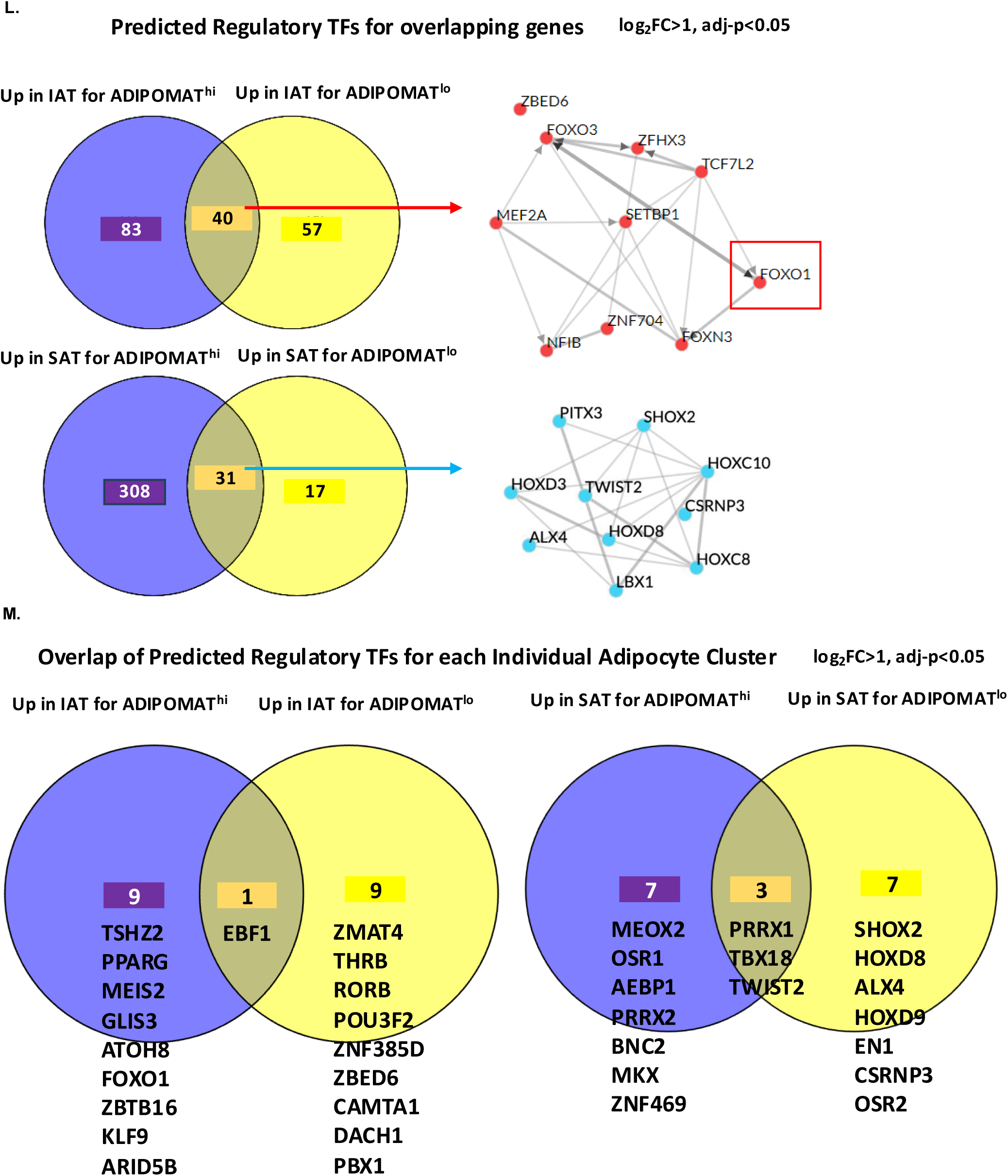
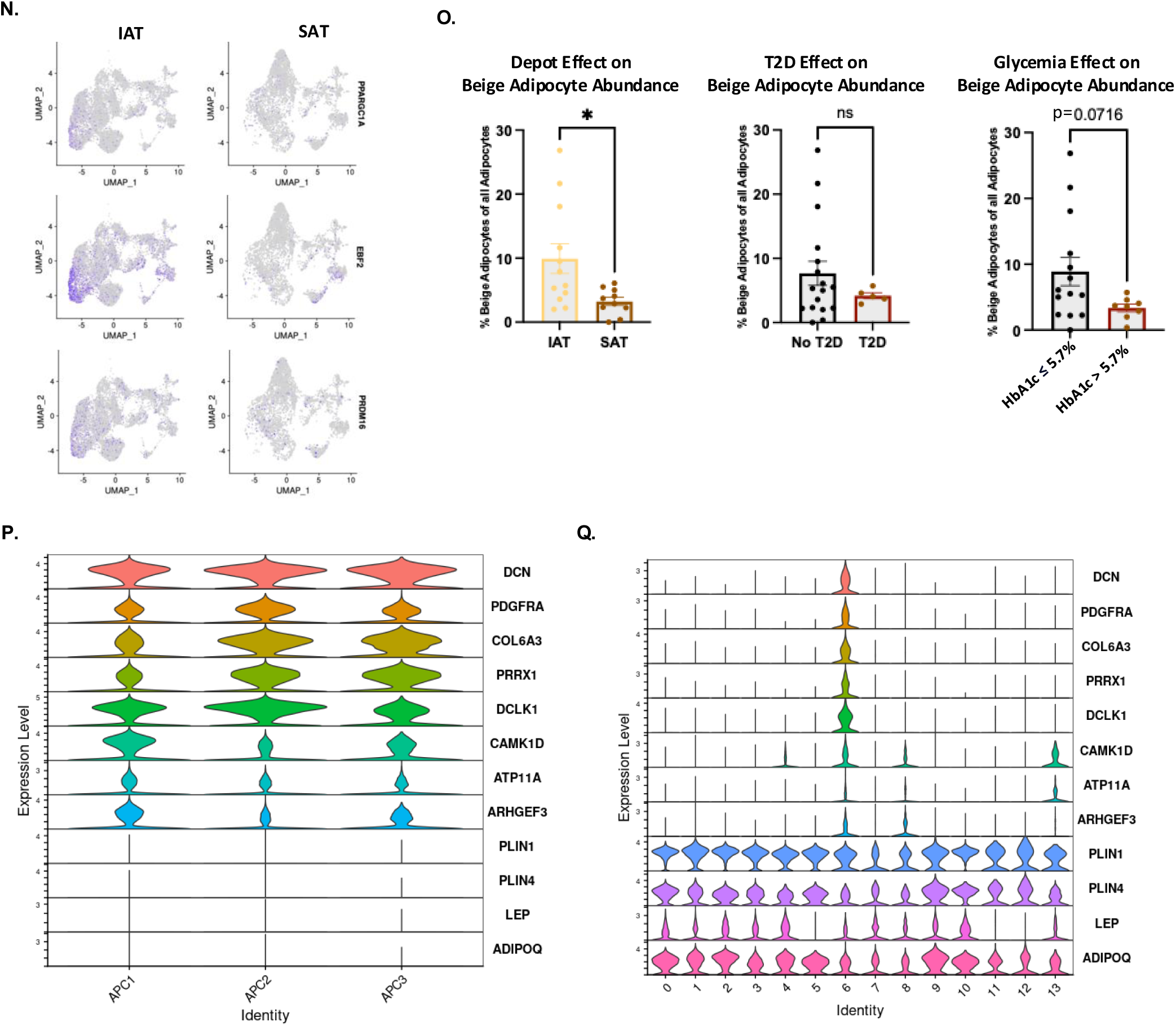
A. Heatmap demonstrating the markers of each distinct APC subtype, B. ENRICHR ontology analysis showing the top 10 enriched pathways based on unsupervised markers of APC1, APC2, and APC3 populations (adjusted-p<0.05, log_2_ fold change > 0.5 as input). Length of the bar signifies −log_10_ p-value C. Venn diagrams of genes upregulated in IAT vs. SAT (left) or SAT vs. IAT (right) for all APC populations and corresponding predicted regulatory transcription factors of the overlapping genes using ChEA3 analysis. D. Venn diagrams of predicted regulatory transcription factors (identified via ChEA3 analysis) for genes that are upregulated in IAT vs. SAT (left) or in SAT vs. IAT (right) for all APC populations. E. Adipogenesis Transcriptomic Score (quantified in VISION) in ADIPOMAT^hi^ and ADIPOMAT^lo^ adipocyte subpopulations within each individual biopsy, F, G. Selected markers of ADIPOMAT^hi^ (F) and ADIPOMAT^lo^ (G), visualized by violin plots. H. Transcriptional signature mapping of our Adipo2 or ADIPOMAT^lo^ to the mature adipocytes from Lazarescu et al.^25^, I. Violin Plots showing expression of ANK2, PTPRB, and PDE4D in our subpopulations of ADIPOMAT^hi^ and ADIPOMAT^lo^ adipocytes and immunofluorescent staining of ANK2, PTPRB, and PDE4D in human adipose tissue samples from Lazarescu et al^25^, J. ENRICHR pathway ontology analysis showing the top 10 enriched pathways for genes differentially expressed between SAT vs. IAT (left) or IAT vs. SAT (right) for ADIPOMAT^hi^ (upper) or ADIPOMAT^lo^ (lower) adipocytes (log_2_FC>0.5, adj-p<0.05). Blue color of the bar signifies significant enrichment for the respective pathway and length of the bar signifies −log_10_ p-value. K. DotPlot and corresponding violin plots showing differentially expressed genes that are upregulated in ADIPOMAT^lo^ vs. ADIPOMAT^hi^ adipocytes within IAT (left) or SAT (right) biopsies, L. Overlap of genes upregulated in IAT vs. SAT (upper) or in SAT vs. IAT (lower) for ADIPOMAT^hi^ and ADIPOMAT^lo^ adipocytes, with corresponding predicted transcription factors (ChEA3 analysis). M. Overlap of predicted regulatory transcription factors (ChEA3 analysis) for genes upregulated in IAT vs. SAT (left panels) or in SAT vs. IAT (right panels) for ADIPOMAT^hi^ and ADIPOMAT^lo^ adipocytes. N. Feature Plot of mature adipocytes showing expression of PPARGC1A, EBF2, and PRDM16, O. Percentage of beige adipocytes (expressed as % of all mature adipocytes) in SAT and IAT biopsies (left), or according to T2D status (middle), or glycemic control (right). Graphs are presented as mean +/− SEM, P, Q. Violin Plots showing expression of DCN, PDGFRA, COL6A3, PRRX1, DCLK1, CAMK1D, ATP11A, ARHGEF3, PLIN1, PLIN4, LEP, and ADIPOQ in the three identified APC populations (P) and in all subpopulations of mature adipocytes (Q).

**Supplementary Figure 3.**
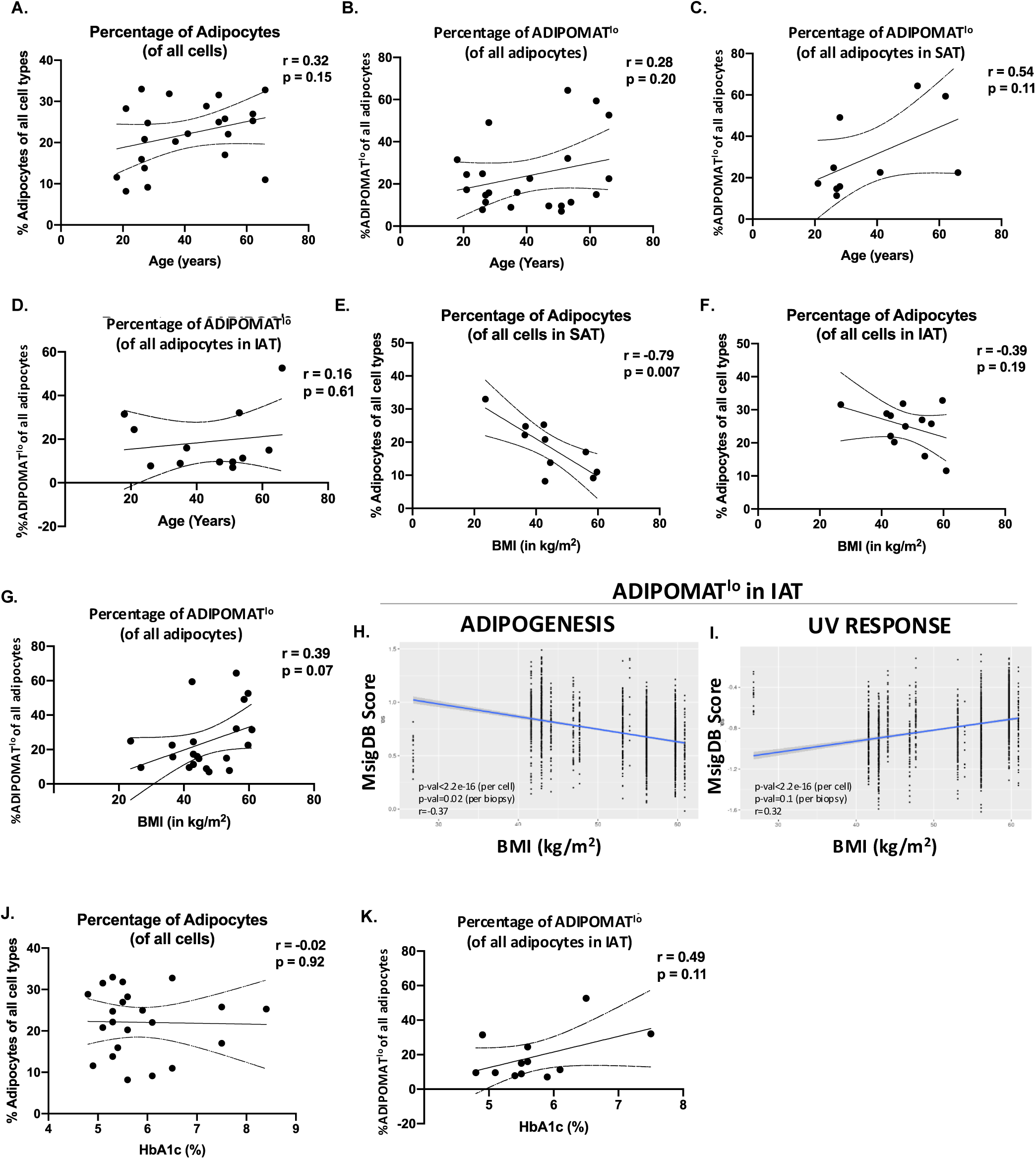
A. Correlation analysis of percentage of adipocytes (expressed as % of all cells) and donor age (years). B. Correlation of percentage of ADIPOMAT^lo^ (expressed as % of all mature adipocytes) and donor age.C. Correlation of percentage of ADIPOMAT^lo^ (expressed as % of all mature adipocytes) in SAT biopsies and donor age. D. Correlation of percentage of ADIPOMAT^lo^ (expressed as % of all mature adipocytes) in IAT biopsies and donor age. E. Correlation of percentage of adipocytes (expressed as % of all cells) in SAT biopsies and donor body mass index (BMI, kg/m^2^). F. Correlation of percentage of adipocytes (expressed as % of all cells) in IAT biopsies and donor BMI. G. Correlation of percentage of ADIPOMAT^lo^ (expressed as % of all mature adipocytes) and donor BMI. H. Correlation of percentage of adipogenesis transcriptomic score (calculated via VISION) in ADIPOMAT^lo^ adipocytes in IAT biopsies and donor BMI. I. Correlation of percentage of UV response (which reflects a pro-inflammatory and pro-fibrotic transcriptomic signature) transcriptomic score (calculated via VISION) in ADIPOMAT^lo^ adipocytes in IAT biopsies and donor BMI. J. Correlation of percentage of adipocytes (expressed as % of all cells) and donor HbA1c (%). K. Correlation of percentage of ADIPOMAT^lo^ (expressed as % of all mature adipocytes) in IAT biopsies and donor HbA1c (%).

**Supplementary Figure 4.**
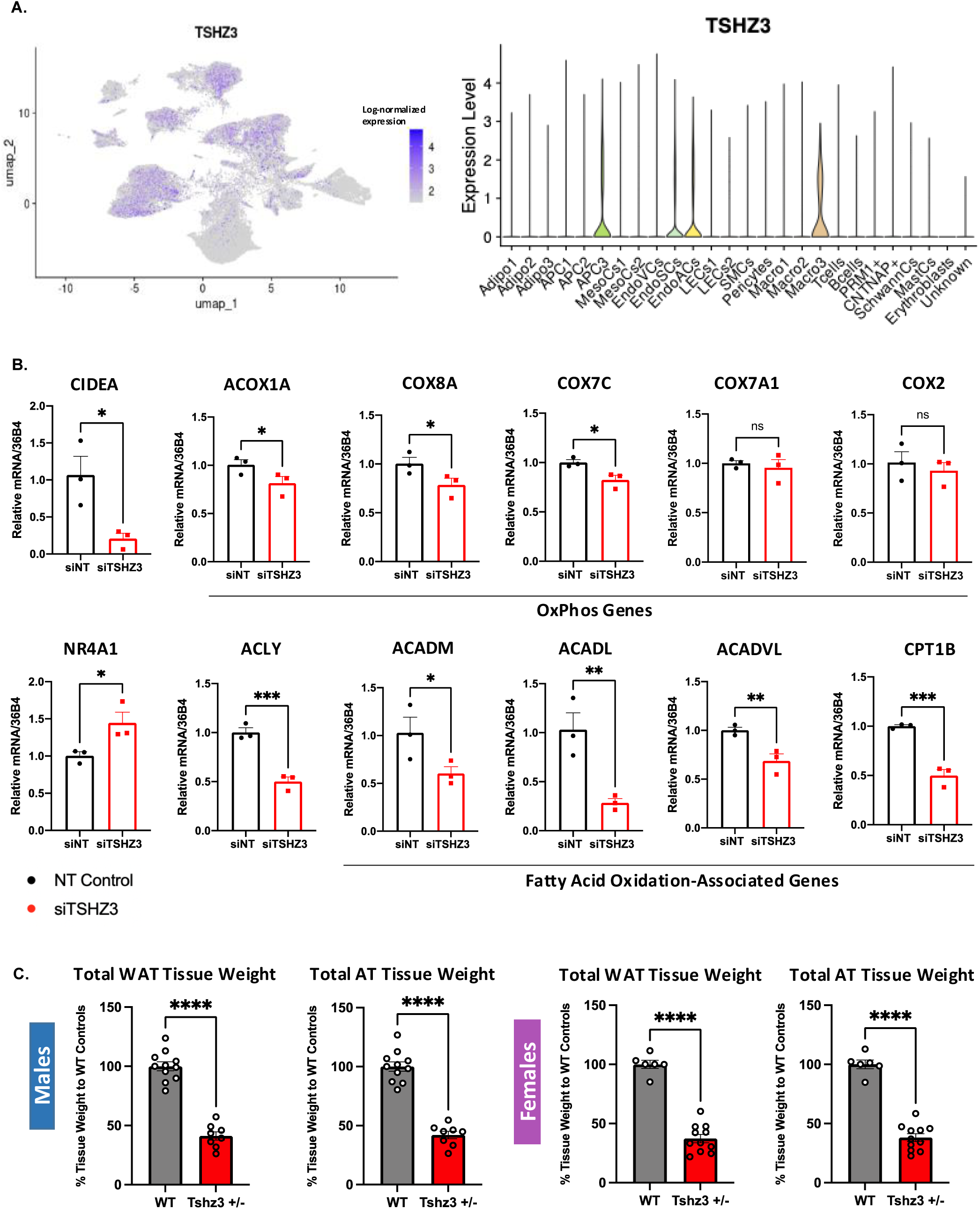
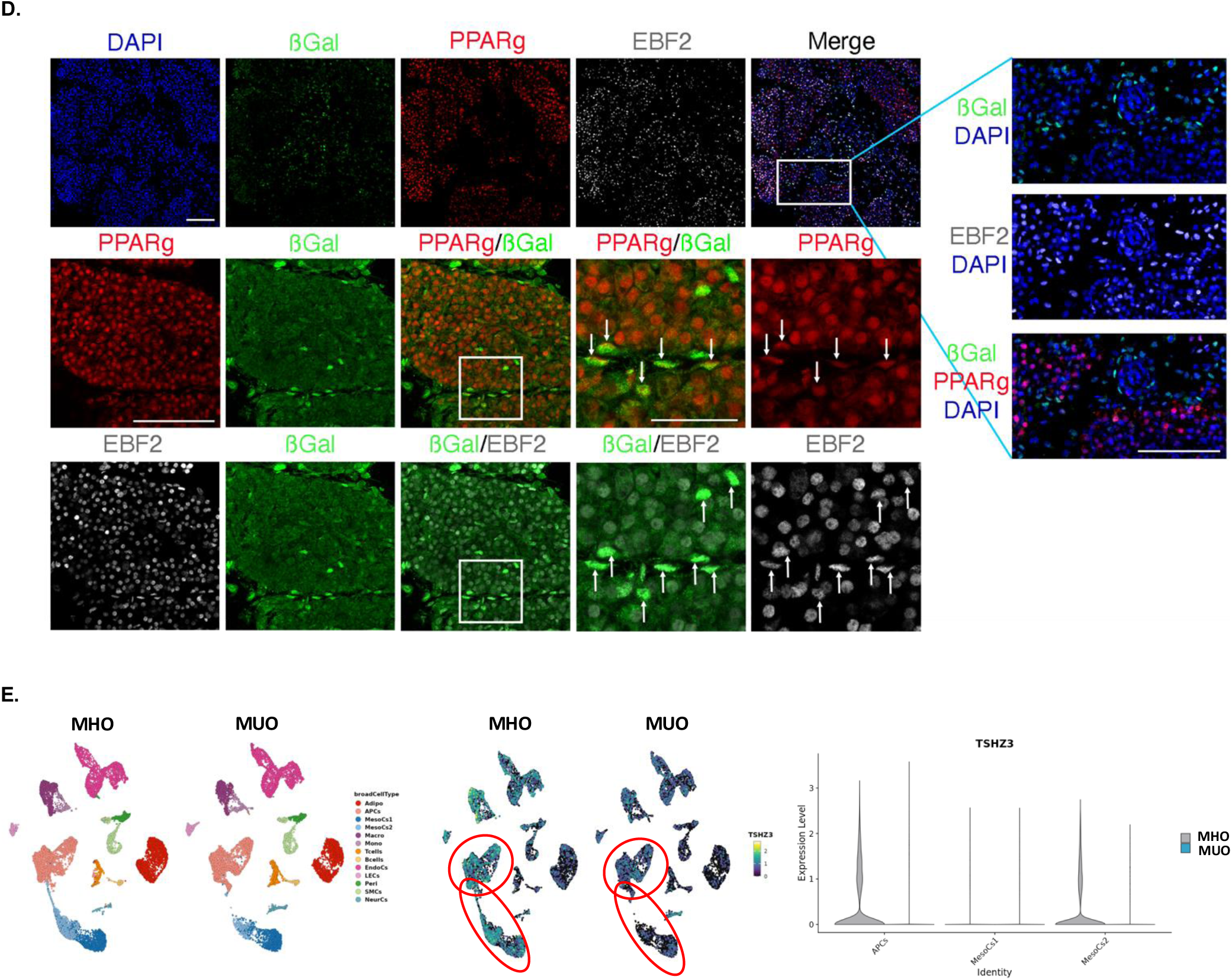
A. Feature Plot (left) and Violin Plot (right) showing expression of TSHZ3 in all identified cell populations. B. Gene expression of CIDEA, ACOX1A, COX8A, COX7C, COX7A1, COX2, NR4A1, ACLY, ACADM, ACADL, ACADVL, and CPT1B (RT-qPCR) in human immortalized mature white adipocytes (day 18 of differentiation) treated with siTSHZ3 or siNT-control on day −2, prior to induction of differentiation. C. Total white adipose tissue (WAT) and total adipose tissue (AT) depot weights in haploinsufficient Tshz3+/lacZ and wild-type control male (left panels) and female (right panels) mice. Graphs are presented as mean +/− SEM. * p<0.05, **p<0.01, ***p<0.001, ****p<0.0001. D. Immunofluorescent imaging of inguinal WAT on embryonic day E18.5 that shows localization of TSHZ3 (B-Gal), EBF2, PPARG, and nuclei (DAPI). E. snRNA-Seq from Reinisch et al.^34^ identified adipose-resident cell populations in MHO and MUO individuals (left); Feature Plot showing TSHZ3 expression in all identified cell populations in MHO and MUO individuals. Red circles signify expression in APCs or mesenchymal-like MesoCs (middle); Violin Plot showing quantification of expression of TSHZ3 in APCs and mesenchymal-like MesoCs in MHO vs. MUO (right).

**Supplementary Figure 5.**
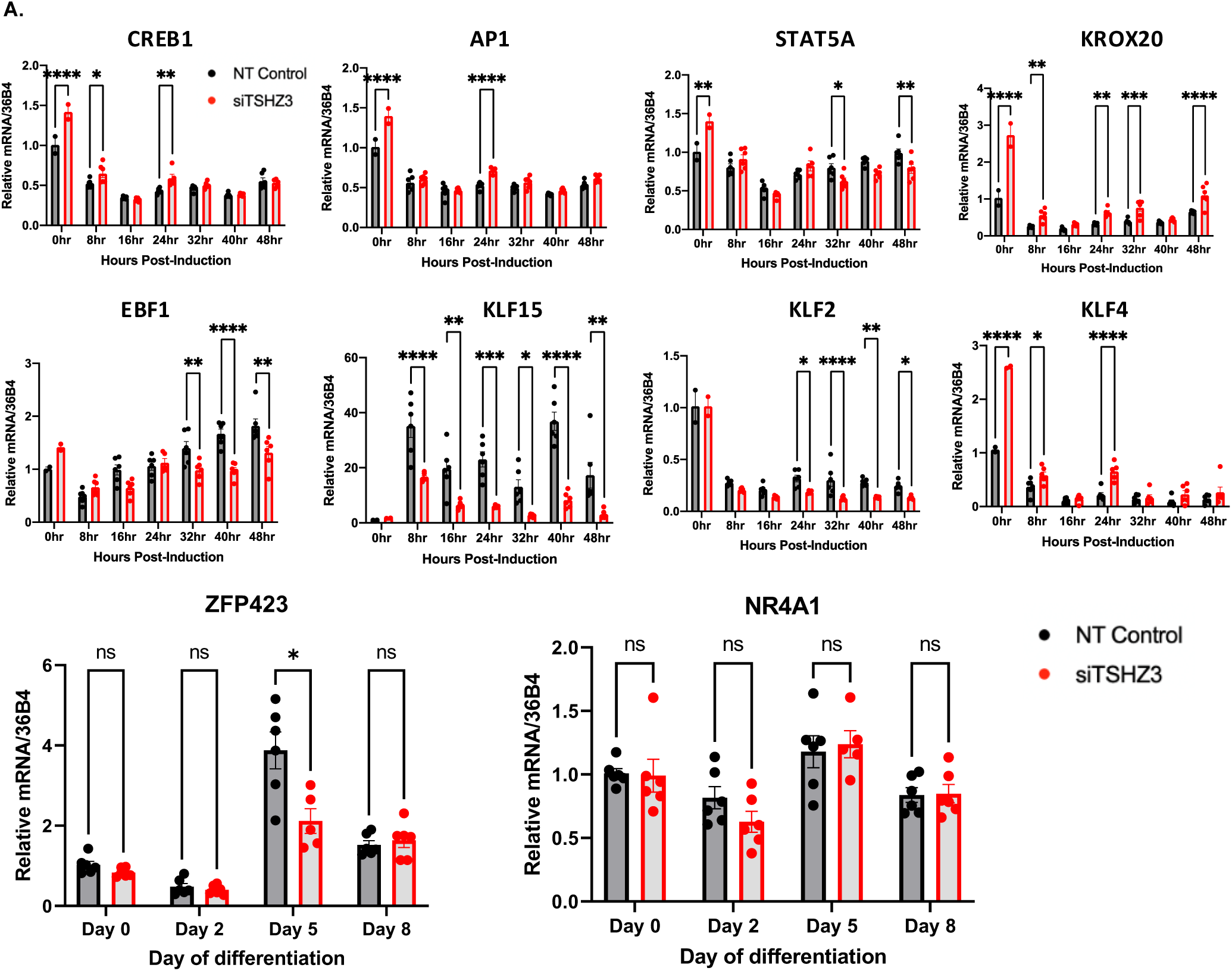
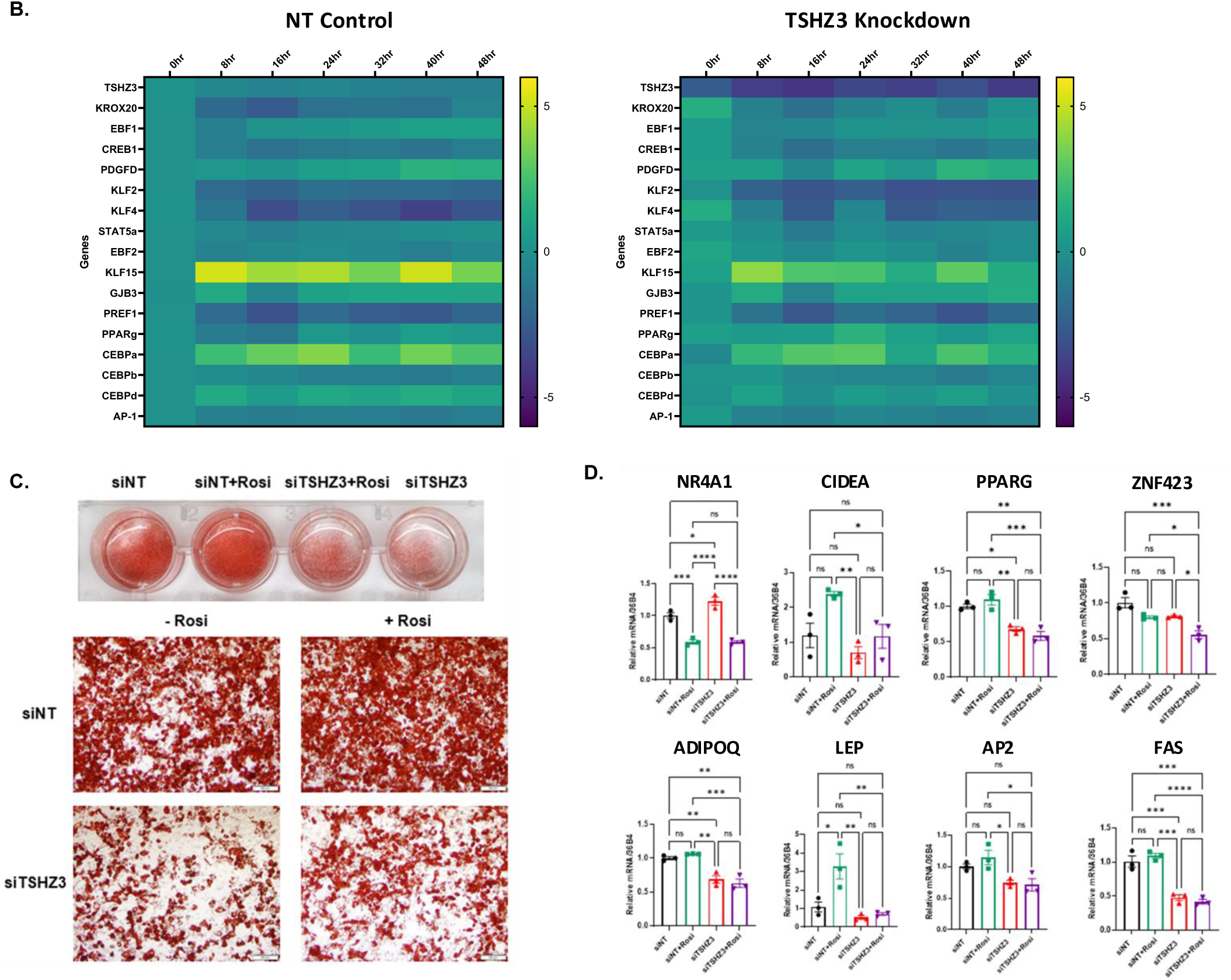
A. Gene expression of CREB1, AP1, STAT5A, KROX20, EBF1, KLF15, KLF2, and KLF4 (RT-qPCR) in human immortalized mature white adipocytes (0-48 hours after differentiation) treated with siTSHZ3 or siNT-control on day −2, prior to induction of differentiation; and gene expression of ZFP423 and NR4A1 (RT-qPCR) in human immortalized mature white adipocytes (days 0-8 of differentiation) treated with siTSHZ3 or siNT-control on day −2. B. Heatmap showing gene expression of TSHZ3, KROX20, EBF1, CREB1, PDGFD, KLF2, KLF4, STAT5A, EBF2, KLF15, GJB3, PREF1, PPARG, CEBPA, CEBPB, CEBPD, and AP1 (RT-qPCR) in human immortalized mature white adipocytes (0-48 hours after differentiation) treated with siTSHZ3 or siNT-control on day −2. Values are normalized to the 0 hr time point of siNT-control cells. C. Impact of siRNA-mediated TSHZ3 vs. nontargeted (siNT) knockdown (on day −2 of differentiation) in human immortalized white adipogenic progenitors in the presence or absence of the PPARG agonist rosiglitazone (added to differentiation medium) on differentiation as assessed by images of Oil Red O-stained cells on day 18 of differentiation. D. Gene expression of NR4A1, CIDEA, PPARG, ZNF423, ADIPOQ, LEP, AP2, and FAS (RT-qPCR) in human immortalized mature white adipocytes treated with siTSHZ3 or siNT-control on day −2, in the presence or absence of rosiglitazone, and assessed at day 18 of differentiation. Graphs are presented as mean +/− SEM. * p<0.05, **p<0.01, ***p<0.001, ****p<0.0001.

**Supplementary Figure 6.**
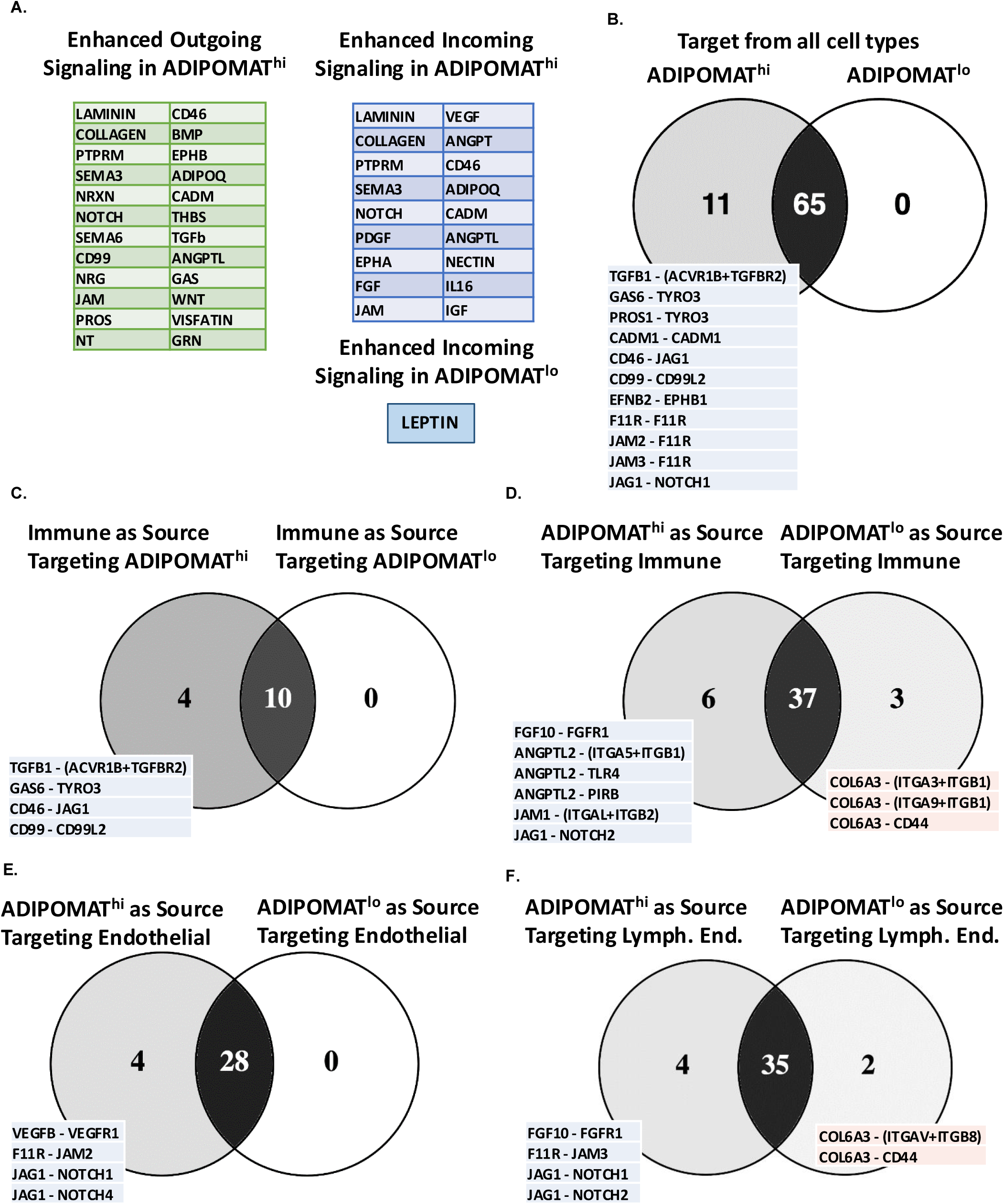
A. Enhanced outgoing signaling (left table) and enhanced incoming signaling (right table) in ADIPOMAThi vs. ADIPOMATlo adipocytes. B. Venn diagram of incoming signaling patterns from all identified cell types to ADIPOMAT^lo^ and/or ADIPOMAT^hi^ adipocytes. C. Venn diagram of incoming signaling patterns from all identified immune cell types to ADIPOMAT^lo^ and/or ADIPOMAT^hi^ adipocytes. D. Venn diagram of outgoing signaling patterns from ADIPOMAT^lo^ and/or ADIPOMAT^hi^ adipocytes to all identified immune cell types. E. Venn diagram of outgoing signaling patterns from ADIPOMAT^lo^ and/or ADIPOMAT^hi^ adipocytes to identified endothelial cells. F. Venn diagram of outgoing signaling patterns from ADIPOMAT^lo^ and/or ADIPOMAT^hi^ adipocytes to identified lymphatic endothelial cells.

